# Comprehensive characterization of the neurogenic and neuroprotective action of a novel TrkB agonist using mouse and human stem cell models of Alzheimer’s Disease

**DOI:** 10.1101/2023.05.08.539797

**Authors:** Despoina Charou, Thanasis Rogdakis, Alessia Latorrata, Maria Valcarcel, Vasileios Papadogiannis, Christina Athanasiou, Alexandros Tsengenes, Maria Anna Papadopoulou, Dimitrios Lypitkas, Matthieu D. Lavigne, Theodora Katsila, Rebecca C. Wade, M. Zameel Cader, Theodora Calogeropoulou, Achille Gravanis, Ioannis Charalampopoulos

## Abstract

Neural stem cell (NSC) proliferation and differentiation in the mammalian brain decreases to minimal levels postnatally. Nevertheless, neurogenic niches persist in the adult cortex and hippocampus in rodents, primates and humans, with adult NSC differentiation sharing key regulatory mechanisms with development. Adult neurogenesis impairments have been linked to Alzheimer’s Disease (AD) pathology. Addressing these impairments is a promising new avenue for therapeutic intervention based on neurogenesis. However, this possibility has been hindered by technical difficulties of using in-vivo models to conduct screens, including working with scarce NSCs in the adult brain and differences between human and mouse models or ethical limitations. In our study, we use a combination of mouse and human stem cell models to circumvent these issues and perform comprehensive characterization of a novel neurogenic compound using *in vitro* screening. Our work focuses on the brain-derived neurotrophic factor (BDNF) pathway, a pivotal neurotrophin in the regulation of neuronal growth and differentiation via its receptor tyrosine receptor kinase B (TrkB). We describe the design, chemical synthesis and biological characterization of ENT-A011, a steroidal dehydroepiandrosterone (DHEA) derivative and BDNF mimetic with neuroprotective and neurogenic actions. The compound is able to increase proliferation of mouse primary adult hippocampal NSCs and embryonic cortical NSCs, in the absence of EGF/FGF, while reducing Amyloid-β (Aβ) induced cell death, acting specifically through TrkB activation. The compound is also able to increase astrocytic gene markers involved in NSC maintenance, protect hippocampal neurons from Αβ toxicity and prevent synapse loss after Aβ treatment. To provide a translational link to human cells, we also used neural progenitor cells (NPCs) differentiated from three human induced pluripotent stem cell lines from healthy and AD donors. Our findings suggest that ENT-A011 successfully induces proliferation and prevents cell death after Aβ toxicity of human NPCs. Additionally, using RNAseq profiling, we demonstrate that the compound acts through a core gene network shared with BDNF. Our work characterizes a novel synthetic BDNF mimetic with potential neurogenic and neuroprotective actions in Alzheimer’s disease via stem cell-based screening, demonstrating the promise of stem cell systems for short-listing competitive candidates for further testing.

## INTRODUCTION

From early embryonic development until early postnatal stages, neural stem cells (NSCs) proliferate, migrate, differentiate and mature to new neurons in a multi-step process known as neurogenesis^1–3^. Neurogenesis is a process that drops sharply postnatally, yet many studies have demonstrated neurogenesis in aged brains of rodents, non-human primates and humans ^4–11^. More specifically, neurogenesis has been detected in the adult mammalian brain in specific areas known as neurogenic niches, namely the subventricular zone (SVZ) of the cortex and the dentate gyrus of the hippocampus^4, 12, 13^. In pathology, dysfunctional adult neurogenesis has been linked to Alzheimer’s disease (AD), a neurodegenerative disease that is the most common cause of dementia, characterized by β-amyloid (Aβ) deposition and neurofibrillary tangle formation. The most affected areas in the AD brain are the cortex and the hippocampus and impaired adult hippocampal neurogenesis (AHN) has been shown to occur in early stages of the development of the disease ^14–16^. Moreover, neural stem cells and progenitor cells from the AD brain have showed reduced proliferation, viability, differentiation, increased senescence when exposed to a harmful microenvironment and decreased interactions with the neurogenic niche^17–21^.

A number of microenvironment processes relating to adult neurogenesis may differ to those involved in embryonic and postnatal neurogenesis, but there are also shared mechanisms at play, such as the role of many trophic factors^22^. Experimental cooption of neurogenic molecular signals *in vitro* has allowed the development of functional central nervous system (CNS) neurons from stem cells derived from adult hippocampus or astrocytes from postnatal hippocampus. Furthermore, the maturation and synaptogenesis mechanisms used by the neural progeny of adult neural stem cells, including factors contributed by neonatal astrocytes, also share similarities with those during development^23^.

Brain-derived neurotrophic factor (BDNF) is a member of the neurotrophin family of trophic factors that binds with high affinity and activates the Tropomyocin receptor kinase B (TrkB), a widely expressed and distributed receptor in the CNS. Many studies have shown the critical role of BDNF and TrkB signaling in adult neurogenesis ^24–26^, while a reduction of BDNF has been reported in the cortex and the hippocampus of AD ^27–30^. BDNF is instrumental in promoting the proliferation, survival, differentiation and functional integration of newborn neurons into hippocampal circuits and the recovery of synaptic degeneration, through the activation of TrkB ^21, 26, 31, 32^. Additionally, BDNF has important roles in the function of the adult neurogenic niche, able to increase and maintain its hippocampal levels through a positive feedback loop. Elevated astrocytic BDNF can increase the efficacy of astrocytes in supporting AHN and countering AD deficits, while exogenous BDNF supplementation combined with independent enhancement of AHN can attenuate cognitive decline in AD ^23, 33–36^. It is of note that several factors, such as exercise or antidepressant drugs induce adult neurogenesis by increasing the levels of endogenous BDNF^33–37^.

Although BDNF has been demonstrated to hold potential in altering AD pathology and counter cognitive decline through its roles in the neurogenic niche and the function of NSCs, its inability to cross the blood-brain barrier (BBB), its poor pharmacokinetic properties and its vulnerability to proteolytic cleavage remain important limitations in using BDNF as a reliable therapeutic agent. This led to the development of small molecules that can activate TrkB signaling and mimic the beneficial properties of BDNF, such as 7,8-dihydroxy flavone (7,8-DHF) and its derivative CF3CN, some of the best studied BDNF mimetics that can prevent neurotoxicity *in vitro* and *in vivo*, enhance learning, prevent memory impairment and promote axonal regeneration ^38–42,43^. Other noticeable examples include GSB-106^44^, LM22A-4 ^45^ that can promote neurite outgrowth and suppress neuronal death in in-vitro neurodegenerative disease models and a derivative of LM22B-10 that is able to increase survival of iPSC-derived cholinergic neurons after exposure to Aβ^46, 47^.In addition, derivatives of DHEA, an endogenous neurosteroid produced by neurons and glia in the Central Nervous System (CNS), that bind to and activate neurotrophin receptors have been explored recently^48^, including TrkA specific activator ENT-A013, TrkA and p75NTR activator BNN27, and BNN20 which activates TrkA, TrkB and p75NTR^49,50^and is able to increase BDNF levels through TrkB signaling and exert dopaminergic neuroprotection ^51–53^.

In the present study, we characterize ENT-A011, a new small molecule mimetic of BDNF that in a structure-specific manner binds to and activates the TrkB receptor and describe its chemical synthesis, potential TrkB binding sites, its metabolic stability and CYP-mediated reaction phenotyping. The compound upregulates genes involved in neurogenic niche maintenance, increases the proliferation, survival and mitochondrial activity of mouse adult and embryonic NSCs and promotes the adult NSC differentiation towards neurons and astrocytes. Moreover, we show ENT-A011 promotes cell survival, protection against synaptic loss and neurite outgrowth in hippocampal and cortical neurons. We then demonstrate ENT-A011 can increase proliferation and counter Aβ-induced cell death in human induced pluripotent stem cell (iPSC) derived NSCs from both control and AD lines. Using RNA sequencing (RNA-seq), we also provide evidence that this compound acts on a defined core gene network shared with BDNF in human NSCs. Overall, we present a small molecule, TrkB specific activator with BDNF-like effects on neurogenesis in rodent and human cells, exerting strong neuroprotective effects against Aβ toxicity, representing a potential lead molecule with high potential for therapeutic applications in AD.

## MATERIALS AND METHODS

### Chemistry

Reactions were run in flame-dried glassware under an atmosphere of argon or nitrogen. All solvents were dried and/or purified according to standard procedures prior to use. Melting points were determined with an Electrothermal Digital Melting Point Apparatus, Cole-Parlmer ET0001/Version 1.0, and were uncorrected. Optical rotations were measured with a P3000 series polarimeter (Krüss Optronic, Hamburg, Germany). NMR spectra were recorded on Varian spectrometers (Varian, Palo Alto, CA, USA). The specific rotation 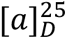 was calculated according to the formula 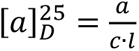. The concentration of the sample is expressed in g/mL. ^1^H NMR spectra were recorded at 300 MHz or 600 MHz, ^13^C NMR spectra were recorded at 75 MHz or 150 MHz and were internally referenced to residual solvent peaks. Chemical shifts are reported in *δ* units, parts per million (ppm) and coupling constants (*J*) are given in Hz. Low-resolution mass spectra were recorded on a LC-MSn Fleet mass spectrometer (Thermo Scientific,Waltham, MA, USA) using MeOH as solvent. High-resolution mass spectra (HRMS) were recorded on UPLC-MSn Orbitrap Velos mass spectrometer (Thermo Scientific). Flash column chromatography (FCC) was performed on silica gel 60 (230–400 mesh, Merck, Darmstadt, Germany) and thin-layer chromatography (TLC) on pre-coated glass plates 60 F254 (0.2 mm, Merck). Spots were visualized with UV light at 254 nm and phosphomolybdic acid stain (PMA, 10% in absolute ethanol). The purity of ENT-A011 and ENT-A012 was determined by high-performance liquid chromatography (HPLC) using Nucelosil 100-5 C18 HD column, 5μm (4.6 x 250 mm), eluting with H_2_O, 0.1% HCOOH – MeOH, 0.1% HCOOH gradient, flow rate 1 mL/min, UV detection at λ = 237 nm. Gradient information: 0.0 – 5.0 min ramped from 5% H_2_O, 0.1% HCOOH – 95% MeOH, 0.1% HCOOH to 100% MeOH, 0.1% HCOOH; 5.0 – 10.0 min held at 100% MeOH, 0.1% HCOOH; 10.0 – 25.0 min returned to 5% H_2_O, 0.1% HCOOH – 95% MeOH, 0.1% HCOOH.

ENT-A011: *t_R_* = 6.32 min, purity = 99.09%. ENT-A012: *t_R_* = 6.32 min, purity = 98.47%. (Supplementary Data, Figures S3 and S4).

### Synthesis of ENT-A011 and ENT-A012

All the detailed chemistry describing the synthesis of compounds ENT-A011 and ENT-A012 is analytically presented in Supplementary Data.

### Metabolic stability

Incubation conditions ensured linear metabolite formation with respect to reaction time and protein concentration (pooled human liver microsome concentration was set at 0.5 mg/mL). To determine the oxidative (CYP-mediated) metabolic stability profile, 1 mM NADPH served as a cofactor. ENT-A011 was tested at 1 μM. Triplicate reactions took place at 37 °C in the presence of negative and positive controls (low vs. rapid clearance). Reactions were terminated after 60 minutes and readouts were recorded by Lionheart FX (BioTek) to determine the residual (%) of time zero (ENT-A011 depletion, Figure S7).

### Isozyme-specific CYP450-metabolism

CYP1A2, CYP2A6, CYP2B6, CYP2C9, CYP2C19, CYP2D6, and CYP3A4 human CYP450 isoenzymes were expressed in Baculosomes®, purchased from Thermo Fisher Scientific (Waltham, MA, USA). All reagents were handled and prepared according to the manufacturer’s protocol. ENT-A011 was tested at 1 μM. On the basis of the kinetic model for each CYP450 isoform in question, CYP450-enzymatic activity was determined in the presence of ΕΝΤ-Α011. Triplicate reactions took place at 20°C, following their initiation via the conversion of NADP+ (10 mM in 100 mM potassium phosphate, pH 8.0) into NADPH by the regeneration system present (glucose-6-phosphate at 333mM and glucose-6-phosphate-dehydrogenase at 30 U/mL in 100 mM potassium phosphate, pH 8.0). Next, the fluorescent substrate was added and immediately (< 2 minutes), signal monitoring over time took place at suitable excitation and emission wavelengths by Lionheart FX (BioTek). CYP450 inhibition (%) was determined based on the reaction rates (fluorescence intensity changes per unit time). In total, n=60 measurements per minute were acquired (t=60 minutes).

% Inhibition= (1-X/A) x 100% where X is the rate observed in the presence of test compound and A is the rate observed in the presence of negative (solvent, DMSO) control (Figure S7).

### Molecular modelling

The crystal structure of the complex of the TrkB immunoglobulin-like domain 5 (D5) homodimer with the mature neurotrophin-4/5 (NT-4/5) homodimer (PDB ID: 1HCF)^54^ was used for modeling studies. The first three residues of the TrkB-D5 are cloning artefacts and thus they were mutated to the wild-type sequence. The structure of NT-4/5 has the following missing residues: Asn 145 – Gly 150 and Gly 207 – Ala 210 in chain A and Gly 81 – Glu 84, Asn 145, Arg 209 and Ala 210 in chain B (residue numbers correspond to the UniProt sequence of human NT-4/5, UniProt code: P34130). The missing residues were built with the AutoModel class of MODELLER v.9.23^55^. The linker corresponding to residues Asn 145 – Gly 150 was built using the structure of human NT-4/5 from PDB ID: 1B98^56^ as a template. The final model was prepared with the Protein Preparation Wizard^57, 58^ of Schrödinger Suite v.2020. Specifically, bond orders were assigned, hydrogens were added, disulfide bonds were created, protonation states were assigned at pH 7 with PROPKA,^59^ and optimization of the H-bond network and restrained energy minimization using the OPLS3e force field^60^ with heavy atoms restricted to a maximum root mean squared deviation (RMSD) from the initial structure of 0.3 Å were performed. For the identification of possible binding sites, the SiteMap tool^61–63^ of the Schrödinger Suite was used. From the predicted binding sites, the corresponding sites 1a and 1b, previously described^64^ as the possible binding sites of the TrkA agonist BNN27, were used for docking studies. Compound ENT-A011 was built in Maestro^65^ and prepared with the LigPrep tool^66^ of the Schrödinger Suite. The compound was docked to the two symmetric sites 1b of the TrkB – NT-4/5 complex using the standard precision (SP) protocol of the Glide tool^67–70^ of the Schrödinger Suite. Grids were generated with Glide to describe the physicochemical properties of the protein pockets. The hydroxyl groups of the serine, threonine and tyrosine residues and the thiol groups of cysteine residues located within the grid box that describes the binding sites were allowed to rotate during the docking calculation in order to capture all possibilities for making hydrogen bonds with the ligands. For site 1b, the Glide SP scores for the two compounds were about –3.5 kcal/mol, indicating modest binding affinity. For site 1a, which is narrower than site 1b, ligand poses with favorable scores could not be obtained with rigid-body docking and therefore, the Induced Fit Docking (IFD) tool^71–74^ of the Schrödinger Suite was used. This utilizes Glide for docking and Prime^75–77^ for protein remodeling, and resulted in docking poses with scores of about –10.5 kcal/mol. In order to do a direct comparison of the docking scores in the two sites, IFD calculations of ENT-A011 in site 1b were also performed giving a docking score of –4.8 kcal/mol. The low scores for both SP and IFD protocols in site 1b are likely due to the higher solvent accessibility of this site compared to site 1a, which is less accessible and more buried at the interface of the receptor with the neurotrophin. Graphical analysis of the docking poses was carried out with VMD v.1.9.3^78^.

### Cell lines

NIH-3T3 cells were stable transfected with human TrkB or TrkC plasmid, provided by Dr C.F.Ibáñez. Cells were cultured in DMEM medium (11965084, Gibco, Grand Island, New York, USA) containing 10% Fetal Bovine Serum (10270106, Gibco, Grand Island, New York, USA), 100 units/ml Penicillin and 0.1mg/ml Streptomycin (15140122, Gibco, Grand Island, New York, USA) at 5% CO_2_ and 37°C.

PC12 and HEK293T cells were sourced from LGC Promochem GmbH (Teddington, UK). PC12 cells were grown in DMEM medium (11965084, Gibco, Grand Island, NY, USA) containing 5% Horse Serum (16050122, Gibco, Grand Island, NY, USA), 10% Fetal Bovine Serum (10270106, Gibco, Grand Island, NY, USA), 100 units/mL Penicillin and 0.1 mg/mL Streptomycin (15140122, Gibco, Grand Island, NY, USA) at 5% CO_2_ and 37°C. HEK293T were grown in DMEM medium (11965084, Gibco, Grand Island, NY, USA) supplemented with 10% Fetal Bovine Serum and they were transfected transiently with the human p75NTR plasmid using lipofectamine.

HEK293 stably expressing kinase NOMAD/tGFP Biosensor (Innoprot), were cultured in DMEM medium (11965084, Gibco, Grand Island, New York, USA) containing 10% Fetal Bovine Serum (10270106, Gibco, Grand Island, New York, USA), 100 units/ml Penicillin and 0.1mg/ml Streptomycin (15140122, Gibco, Grand Island, New York, USA) and 1X MEM Non-Essential Amino Acids Solution (11140050, Gibco, Grand Island, New York, USA) at 5% CO2 and 37oC. Cells were transiently transfected with cDNA plasmid encoding TrkB using calcium phosphate transfection method.

### Animals

All primary cells were isolated from C57BL/6J mice (Charles River Laboratories). Mice had free access to food and water and were housed under a 12-hour light-dark cycle. For cell isolation, mice were deeply anesthetized with ketamine/xylazine and decapitated. All procedures were performed under the approval of Veterinary Directorate of Prefecture of Heraklion (Crete) and carried out in compliance with Greek Government guidelines and the guidelines of FORTH ethics committee and were performed in accordance with approved protocols from the Federation of European Laboratory Animal Science Associations (FELASA) and Use of Laboratory animals. License number: EL91-BIOexp-02, Approval Code: 360667, Approval Date: 29/11/2021 (active for 3 years).

### Primary astrocytes

Mixed glial cultures were isolated from the cortex of C57B/6 pups at post-natal day 2 (P2). Cells were plated in medium containing high-glucose DMEM, 200 U/ml penicillin, 200 µg/ml streptomycin and 10% fetal bovine serum (FBS). When cells reach 100% confluency (7–8 days), the anti-mitotic agent Ara-C was added in the media at a final concentration of 10 µM, for 3 to 4 days to target the highly proliferative microglial cells. Ara-C was removed and primary astrocytes (97% purity) were cultured at 5% CO_2_ and 37°C.

### Primary neural stem cell cultures

Primary adult hippocampal Neural Stem Cells (NSCs) were isolated from mice at postnatal day 7 and cultured with DMEM/F-12 (Dulbecco’s Modified Eagle Medium/Nutrient Mixture F-12) medium (ThermoFischer Scientific, cat# 11320033) containing 2% B27 wihtout Vitamin A (ThermoFischer Scientific, cat# cat#12587010), 50 mg/mL Primocin (Invitrogen, cat# ant-pm-1 500mg), 20 μg/mL FGF (233-FB-025, R&D, cat#), 20 μg/μl EGF (236-EG-200, R&D) and 50mg/mL Heparin. For all the experiments cells were plated on PDL/laminin and the assays were performed after 24hrs.

Primary Embryonic Cortical Neural Stem Cells (NSCs) were isolated from E13.5 mice and cultured with DMEM/F-12 (Dulbecco’s Modified Eagle Medium/Nutrient Mixture F-12) medium (11320033, ThermoFischer Scientific) containing 2% B27 without Vitamin A (12587010, ThermoFischer Scientific), 50 mg/mL Primocin (ant-pm-1 500mg, Invivogen), 20 μg/mL FGF (233-FB-025, R&D) and 20 μg/μl EGF (236-EG-200, R&D). For all the experiments cells were plated on PDL/laminin and the assays were performed after 24hrs.

### Primary Hippocampal neurons

E17.5 mouse embryos were used to isolate primary hippocampal neurons, which were then grown in Neurobasal medium, containing 2% B27, 1% PenStrep, 1X GlutaMax, and 10mM HEPES. Cells were kept at 37°C and 5% CO_2_ in a humified incubator. Cells were cultured for 16–18 days before being exposed for 48 hours to ENT-A011 and 5 uM oligomeric A-beta for the TUNEL assay. Following the manufacturer’s instructions, cells were subsequently fixed with 4% paraformaldehyde (PFA) and labelled with TUNEL (Roche, cat# 11684795910). Cells were then immunostained (1:2000, Biolegend, cat# 801201) against Tuj1 and imaged using confocal microscopy. Similarly, c ells were cultured for 16–18 days before being treated for 4 hours with ENT-A011 and 5 uM oligomeric A-beta for the synaptic plasticity assessment. Following that, cells were fixed with 4% PFA and immunostained against Tuj1 and Synaptophysin (1:1000, Invitrogen, cat# PA1-1043). Images acquired using confocal microscopy and total area of synaptophysin positive puncta was measured and normalized to Tuj1 total area.

### Primary cortical neurons

Primary cultures of cortical neurons were prepared from the cerebral cortices of Sprague-Dawley rat foetuses at embryonic day 18. Brains were removed and freed from the meninges, and the tissues were then dissected under a binocular microscope. Neurons were dispersed with trypsin 0.2% and DNase I 0.04% for 10 min at 37°C, and plated on poly-L-lysine coated 96-well plates at a density number of 30.000 cells per well with neurobasal medium supplemented with B27 for 8 days at 37°C in a humidified 5% CO2 atmosphere. At day 8, cells were pre-treated with BDNF (500 ng/ml) and ENT-A011 (1 µM) one hour before being subjected to a glutamate excitotoxicity condition (100 µM 15 min). After glutamate exposure, medium was replaced and cells were incubated for an additional 24 h for neurite outgrowth measurement.

### Human induced-Pluripotent Stem Cells (hiPSCs)

hiPSC lines were kindly provided by Dr Z. Cader (MTA) and they were maintained in mTeSR (85850, STEMCELL Technologies, Vancouver, British Columbia, Canada) media on matrigel substrate (354277, Corning, New York, USA) and mechanically passaged using 0.5mM EDTA (15575020, ThermoFischer Scientific, Rockford, USA). Differentiation was initiated with 10 μM SB431542 (616464, Sigma, Burlington, MA, USA) and 1 μM dorsomorphin (ab120843, Abcam, Cambridge, UK) following published protocols (Shi et al., 2012)^79^. Further patterning was performed in N2B27 media with retinoic acid. Cells were plated at day 18 on laminin (L2020, Sigma, Burlington, MA, USA) coated 96 well plates for the Celltox and the MTT assays, 24well plates for the proliferation assay and 12well plates for the RNA sequencing.

### Immunoprecipitation and Immunoblotting

Cells were plated at 70-80% confluency. Next day, they were deprived from serum for 16hrs and subsequently treated with 500ng/ml BDNF or 1μM of compound ENT-A011 for 20minutes. Cells were then lysed in Pierce™ IP Lysis Buffer (87788, ThermoFischer Scientific, Rockford, USA) containing proteases (539138, Calbiochem, Darmstadt, Germany) and phosphatases inhibitors (524629, Calbiochem, Darmstadt, Germany). Lysates were then immunoprecipitated overnight at 4°C with TrkA antibody (1:100, 06-574, Sigma-Aldrich, St. Louis, MO, USA) or p75^NTR^ antibody (1:100, ab6172, Abcam, Cambridge, UK) followed by 4hrs incubation with protein G-plus agarose beads (sc-2002, Santa Cruz Biotechnology, California, USA). Beads were then collected, washed 3X with lysis buffer, resuspended in SDS loading buffer and subjected to Western Blot against phosphorylated Tyrosine (1:1000, BAM1676, R&D systems, Mineapolis, USA) or Traf6 antibody (1:2000, ab33915, Abcam, Cambridge, UK). Whole cell lysates were subjected to Western Blot against TrkA (1:1000, 06-574, Sigma-Aldrich, St. Louis, MO, USA), phosphorylated TrkB (1:1000, ABN1381, Sigma-Aldrich, St. Louis, MO, USA), phosphorylated TrkC (1:1000, STJ90960, St John’s Laboratory, London, UK), TrkB (1:1000, 07-225-I, Sigma-Aldrich, St. Louis, MO, USA), TrkC (1:1000, C44H5, Cell Signalling Technology, Danvers, MA, USA), p75^NTR^ (1:1000, 839701, Biolegend, San Diego, USA), Traf6 (1:2000, ab33915, Abcam, Cambridge, UK) phosphorylated Akt (1:1000, 9721S, Cell Signalling Technology, Danvers, MA, USA) and total Akt (1:1000, 4691S Cell Signalling Technology, Danvers, MA, USA).

### Quantitative RT-PCR (QPCR)

Total RNA was extracted from cells using TRIzol Reagent (15596026, Thermo Fisher, Waltham, MA, USA), and cDNA was synthesized using the High-Capacity cDNA Reverse Transcription kit (4368814, Thermo Fisher, Waltham, MA, USA) according to the supplier protocols. For qPCR experiments run with SYBR green dye, for 20seconds at 95°C, followed by 40 cycles of 95°C for 3seconds and 60°C for 30seconds on a StepOne Real-Time PCR System (Thermo Fisher Scientific, Waltham, MA, USA. Β-Actin was used as a housekeeping gene to normalize the genes expression levels. Data were collected and analyzed using the StepOne Software v2.3 (Thermo Fischer Scientific, Waltham, MA, USA). Mouse primer sequences used are listed in Supplementary Table.

### CellTox assay

CellTox assay (G8742, Promega, Leiden, Belgium) was used to assess survival of NIH-3T3 TrkB or PC12 cells under serum deprivation conditions, survival of primary hippocampal or embryonic cortical neural stem cells as well as human neural progenitor cells treated with 10μM oligomeric Aβ1-42. Cells were plated in 96-well plates, NIH-3T3 TrkB cells were starved from serum for 24hrs and subsequently treated with BDNF (500ng/ml) or ENT-A011 (1μM) in the presence or absence of TrkB inhibitor ANA-12 at 100μM (SML0209, Sigma-Aldrich, Burlington, MA, USA) for 24hrs. Cells were treated with 10μM oligomeric Aβ1-42 and BDNF (500ng/ml) or ENT-A011 (1μM) for 24hrs and the same conditions with ANA-12 (100μM). CellTox assay reagents and Hoescht (1:10,000, H3570, Invitrogen, Massachusetts, USA) were added with the treatments and at the end of the 24h treatments, cells were imaged with a Zeiss AXIO Vert A1 fluorescent microscope. CellTox positive cells were normalised to total number of cells for each image.

### MTT assay

Following Celltox assay for the primary hippocampal or embryonic cortical neural stem cells, media was removed and the cells were washed with PBS before adding the MTT solution (M2128, Sigma-Aldrich, Burlington, MA, USA), final concentration 0.5mg/ml for 4hours at 37°C, 5% CO_2_. Supernatant was removed and DMSO-isopropanol solution (1:1 ratio) was added to the wells followed by incubation at room temperature for 15minutes and at 4^ο^ C for another 15minutes. Absorbance was measured at 545nm with a reference at 630nm.

### Nomad biosensor assay

Transiently NOMAD/tGFP TrkB-HEK293 cells were plated in 96-well plate and were treated with ENT-A011 at increasing concentrations (100nM, 300nM, 1μM, 3μM, 10μM and 300μM) for a dose response curve. Fluorescence was measured in a plate reader at 485/528 (ex/em) for NOMAD/GFP and at 340/485 (ex/em) for Hoechst 33342. Data were normalized against negative control and nuclei count.

### LPS assay

Primary astrocytes were treated with the ENT-A011 (1μM) and treatments were repeated after 24hours, LPS was added after 2 hours at 100ng/ml. Cells were lysed with Trizol after 4hours.

### Proliferation assay

Deprivation from EGF/FGF for 3hrs was followed by treatment with EGF/FGF (20ng/ml), BDNF (500ng/ml) or ENT-A011 (1μM) for 24hrs for mouse primary neural stem cells. Cells were also treated with 10μM oligomeric Aβ1-42 and BDNF (500ng/ml) or ENT-A011 (1μM) for 24hrs. Human neural progenitor cells plated at day 25, at the following day cells were treated with BDNF or ENT-A011 and treatments were repeated after 24 hours. BRDU was added to the cells 6hrs prior the end of the treatment. Cells were fixed with 4% PFA and immunostained for BRDU (1:200, ThermoFischer Scientific cat#B35128) and Nestin (1:1000, R&D cat#NB100-1604) and imaged with a Zeiss AXIO Vert A1 fluorescent microscope.

### Cell differentiation assay

Primary hippocampal neural stem cells were plated at confluency of 100,000 cells/ml. Following 3 days of plating, fresh medium was added to the cells without EGF/FGF, with or without BDNF (500ng/ml) or ENT-A011 (1μM). Medium changes were performed every other day and the cells were lysed with Trizol and samples were collected or fixed with 4% PFA after 10 days of culture.

### Isolation of RNA and 3′ RNA sequencing

Human neural progenitor cells were plated at day 30 and treatments with BDNF (500ng/ml) or ENT-A011 (1μM) were performed every two days for a total of 10 days. At day 40, total RNA from biological triplicates was extracted using Trizol reagent (Thermo Scientific) as per the manufacturer’s protocol. The quantity and quality of extracted RNA samples were analyzed using RNA 6000 Nano kit on a bioanalyzer (Agilent). RNA samples with RNA integrity number (RIN)>7 were used for library construction using the 3′ mRNA-Seq Library Prep Kit FWD for Illumina (QuantSeq-LEXOGEN) as per the manufacturer’s instructions. Amplification was controlled by qPCR for obtaining optimal unbiased libraries across samples by assessing the number of cycles (14) required for amplification of the library. DNA High Sensitivity Kit for bioanalyzer was used to assess the quantity and quality of libraries, according to the manufacturer’s instructions (Agilent). Libraries were multiplexed and sequenced on an Illumina Nextseq 500 at the genomics facility of IMBB FORTH according to the manufacturer’s instructions and the number of reads obtained are listed in the Supplementary Table 1.

### Differential Expression Analysis (DEA) and Gene Ontology (GO) enrichment analysis

The quality of the FASTQ files was assessed with the FastQC software^80^. Reads were aligned to the human (hg38) genome with the Hisat2 aligner^81^ (hisat2 –p 4 –q –x $reference_genome –U $read_file –S $outFile ––no-spliced-alignment). Htseq-count^82^ was utilized to summarize reads at the gene level (htseq-count $align_file $genes_gtf > $outFile). Differential expression analysis (DEA) was conducted by running EdgeR^83, 84^ via SARTools 1.5.0^85^. For each comparison, ENT-A011 and BDNF treated APOE4 human Cortical Neural Progenitor Cells (NSCs) compared to untreated control cells, differentially expressed genes (DEGs, up and down-regulated) were defined by applying the following threshold *p*-adj<0.1 and were considered statistically significant. Heatmaps were created in R with the Pheatmap R package. GO analysis was run on the webtool Metascape (http://metascape.org)^86^. Results are listed in Supplementary Table 2 and representative GO terms were selected and summarized in Figure 7.

### Preparation of Aβ oligomers

A-beta (1-42) peptide was obtained from AnaSpec (cat# AS-20276, AnaSpec, California, USA) and prepared in accordance with the directions of the manufacturer. The peptide was diluted in DMEM at the appropriate concentration and left for 24 hours at 37°C. Following a 5’ at 15,000g centrifugation, oligomeric A-beta was derived from the supernatant and cells were treated.

### Neurite outgrowth

Rat cortical neurons were pre-treated for 1 hour with BDNF (200 ng/ml) or the compound ENT-A011 (1μM) 24 hours before being subjected to a glutamate excitotoxicity condition where cells were incubated with 100 μM glutamate during 15 min in medium without B-27 component. After glutamate exposure, medium was replaced with neurobasal medium with B27 factor for additional 24 h.

Beta-III tubulin staining was determined by immunocytochemistry. Once cells were stained and image with fluorescent dyes, cells were washed with PBS and fixed with 4% PFA for 15 minutes. After the fixation step the samples were washed three times with PBS and permeabilized with PBS + 0.3% Triton for 10 minutes. The samples were then blocked with PBS and Bovine Serum Albumin (BSA) for 30 minutes and finally anti-tubulin III antibody were added at 1/1000 in PBS and 0.5% BSA for 60 minutes at room temperature. After three washing steps, the secondary antibodies Alexa 633 were added at 1/100 for 60 minutes to react against the primary antibody. The samples were then washed three times and measured in the Pathway 855 automated fluorescent microscope. Neurite average branch per neuron was measured in order to investigate the neurite outgrowth.

## RESULTS

### Chemistry

The synthesis of ENT-A011 and ENT-A012 is reported in Scheme 1 (Supplementary Data) and involves seven high-yielding steps starting from DHEA. Thus, Horner-Wadsworth-Emmons reaction of DHEA with triethylphosphonoacetate using sodium ethoxide (EtONa) as base gave the (*E*)*-α,β*-unsaturated ester 1 in 96% yield ^87, 88^, which was, in turn, reacted with tert-butyldimethylsilyl chloride (TBSCl) to afford the tert-butyldimethylsilyl-protected alcohol 2 in 93% yield. Selective reduction of the ester group in 2 using diisobutylaluminum hydride (DIBAL-H) gave the allylic alcohol 3 in quantitative yield, which was subjected to a Simmons–Smith cyclopropanation reaction in the presence of diiodomethane and diethylzinc to yield the (17*S*,20*S*)*-*cyclopropyl derivative 4 in 60% yield after purification^89, 90^. Compound 4 was subsequently oxidized with Dess–Martin periodinane (DMP) in dichloromethane (DCM) to afford the corresponding aldehyde 5 in 84% yield. Horner-Wadsworth-Emmons reaction of aldehyde 5 with diethyl(cyanomethyl)phosphonate in the presence of NaH afforded the *α,β*-unsaturated nitrile 6, as a mixture of *E,Z* geometrical isomers 6-*E* and 6-*Z* in 60:40 ratio, respectively ^91^. Compounds 6-*E* and 6-*Z* were easily separated by flash column chromatography (FCC) and were deprotected separately using HF. Pyridine complex in dry CH_2_Cl_2_ to yield the final compounds ENT-A011 and ENT-A012, respectively, in quantitative yield. The geometry of the double bond in compounds ENT-A011 and ENT-A012 was confirmed by the corresponding coupling constants of the two vinylic protons of the unsaturated cyano functionality. In particular, in ENT-A011 they resonate at 5.31 ppm (d, *J* = 16.0 Hz, 1H) and 6.30 ppm (dd, *J* = 16.0, 10.2 Hz, 1H) and the coupling constant is equal to 16.0 Hz indicating the *trans* stereochemistry of the double bond (Supplementary Data, Figure S1). In addition, the *cis* stereochemistry in ENT-A012 is confirmed by the coupling constant (J) of the two vinylic protons being equal to 10.8 Hz [5.19 ppm (d, *J* = 10.8 Hz, 1H) and 6.03 ppm (t, *J* = 10.8 Hz, 1H)] (Supplementary Data, Figure S2).

### ENT-A011 activates TrkB and downstream signalling, rescuing NIH-3T3-TrkB cells from serum deprivation-induced cell death through TrkB

Our first aim was to investigate if the new DHEA derivatives are able to mimic the role of BDNF. Thus, their ability to activate the high affinity receptor of BDNF, TrkB, and its downstream signalling, was tested in appropriate cell lines. The NIH-3T3-TrkB stably expressing cell line was used as screening platform and treatments were carried out with BDNF or ENT-A011 / ENT-A012 for 20 minutes followed by Western Blot analysis to measure phosphorylated and total TrkB receptor’s isoforms, as well as phosphorylated and total Akt, as the pro-survival downstream target. While ENT-A012 was unable to activate the receptor (Figure S5), ENT-A011 induced TrkB and Akt phosphorylation at similar levels to BDNF treatment, (Figure 1A). We observed a 1.84-fold change (FC) in TrkB phosphorylation levels upon ENT-A011 administration, a 1.87 FC after BDNF treatment compared to the untreated control, a 2.28 FC in Akt phosphorylation after ENT-A011 and a 2.18 FC after BDNF treatment compared to the control.

**Figure 1.**
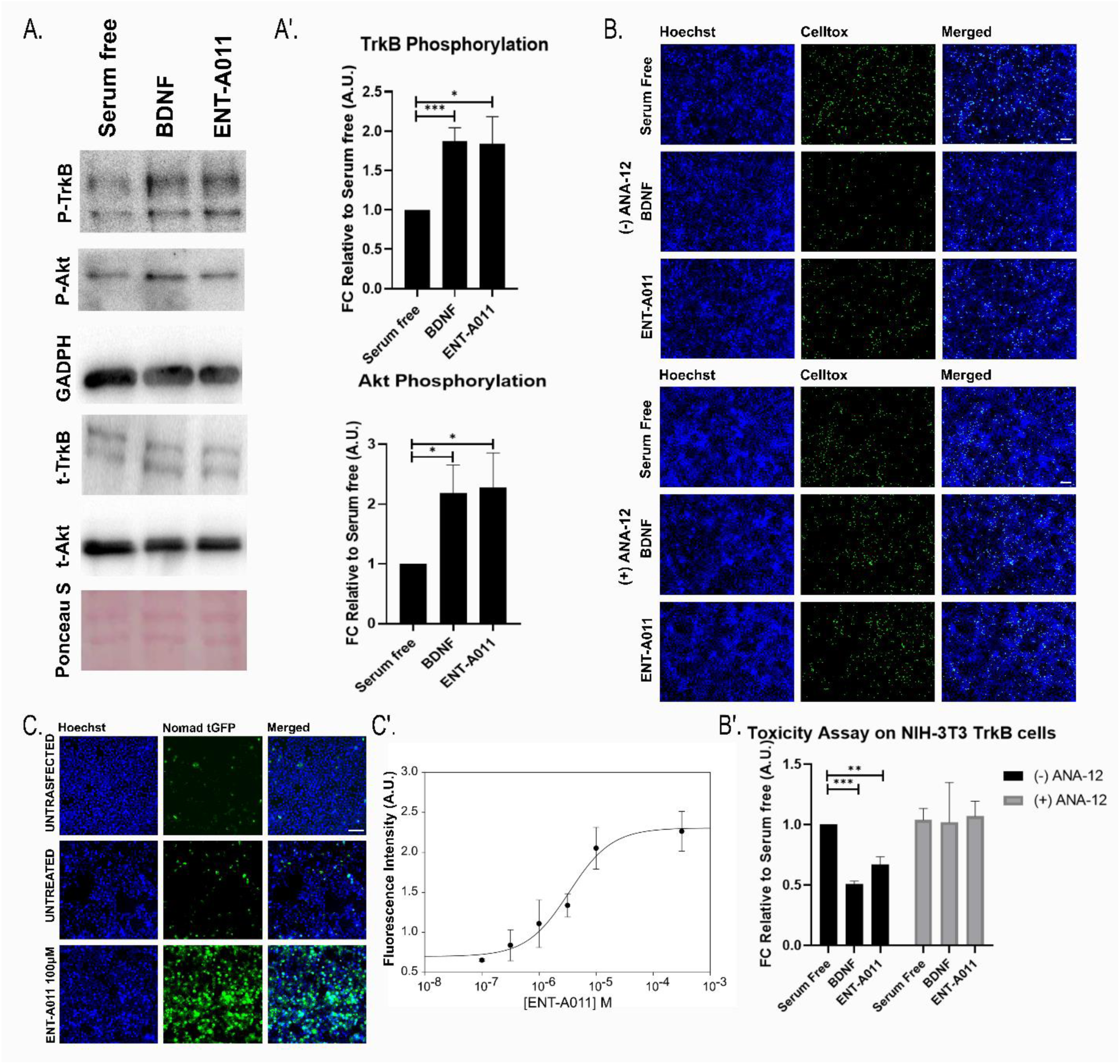
Screening of ENT-A011. ENT-A011 induces TrkB and its downstream target Akt phosphorylation after 20 minutes treatment in NIH-3T3 TrkB stable expressed cells. Representative blots (A) and quantification (A’) of 8 independent experiments are shown. Error bars represent S.E.M., Student’s t-test against Control; * p<0.05; *** p<0.001. B, B’. ENT-A011 protects NIH-3T3 TrkB cells from cell death caused by serum deprivation. Representative images (B) and quantification of Toxicity Assay (B’) on NIH-3T3 TrkB cells after treatment with BDNF or compound ENT-A011 for 24h with or without TrkB inhibitor, ANA-12. N=9-10, error bars represent S.E.M., Student’s t-test against Control; *<0.05, **<0.01, ***<0.001. C, C’. Nomad biosensor activation via TrkB phosphorylation in HEK 293 TrkB transiently transfected cells after ENT-A011 treatment. Receptor activation leads to increase in fluorescence levels. Representative images (C) and dose response curve for different compound concentrations (C’) are shown. Scalebars = 100μm

We then assessed the ability of the compound in preventing cell death. NIH-3T3-TrkB cells underwent serum deprivation for 24 hours and were subsequently subjected to treatment with BDNF or ENT-A011 for another 24 hours. Celltox reagent and Hoescht were used to identify apoptotic cells, in proportion to the total cells in culture. ENT-A011 reduced cell death levels by 33% compared to the untreated control (FC in cell death levels equal to 0,67). We confirmed that this effect is mediated through TrkB receptor activation using the selective TrkB inhibitor, ANA-12, which abrogated the rescuing effect of both ENT-A011 or BDNF (Figure 1B, B’).

Focusing on ENT-A011 for further testing, we also used a NOMAD biosensor cell line, transfected with the TrkB receptor plasmid. In this system, the structural folding of the biosensor changes upon TrkB receptor phosphorylation, leading to an increase in fluorescent readout and allowing for direct receptor activation quantification. Fluorescence increased proportionately with increasing compound concentrations with the IC_50_ calculated at 10.64μM and no toxicity effects observed (Figure 1C, C’).

To gain a structural understanding of the interaction of ENT-A011 with the TrkB receptor, we probed the binding mode of ENT-A011 to TrkB with molecular docking studies. The results showed that ENT-A011 can interact with TrkB similarly to the way the previously identified TrkA agonists BNN27 and ENT-A013 interact with TrkA. Specifically, ENT-A011 can bind in the two pockets (Figure S6) that exist at the interface of the extracellular domain of the TrkB receptor with the neurotrophin and can thus bridge the interactions between neurotrophin and receptor as a possible mode of action. Further information about the computational results can be found in the supplementary information.

### ENT-A011 increases TrkB and Akt phosphorylation, inducing Bdnf, Wnt and Creb gene expression in primary astrocytes

Following results in the TrkB transfected cells, we then used primary astrocytes, as an endogenously TrkB expressing neuronal population, to assess the effects of the compound in a primary cell line that is dependent upon TrkB signalling. ENT-A011 successfully increased levels of phosphorylation of the TrkB receptors as well as the downstream target Akt compared to the untreated control (Figure 2A, A’), with comparable effect to BDNF. We also used RT-qPCR to assess the effect of TrkB activation after ENT-A011 treatment on the expression of key genes involved in the TrkB signalling pathway in astrocytes. In agreement with its ability to activate TrkB receptor, the compound led to an increase in the expression of *Bdnf*, *Wnt* and *Creb*, following astrocyte activation by Lipopolysaccharide (LPS) (Figure 2B).

**Figure 2.**
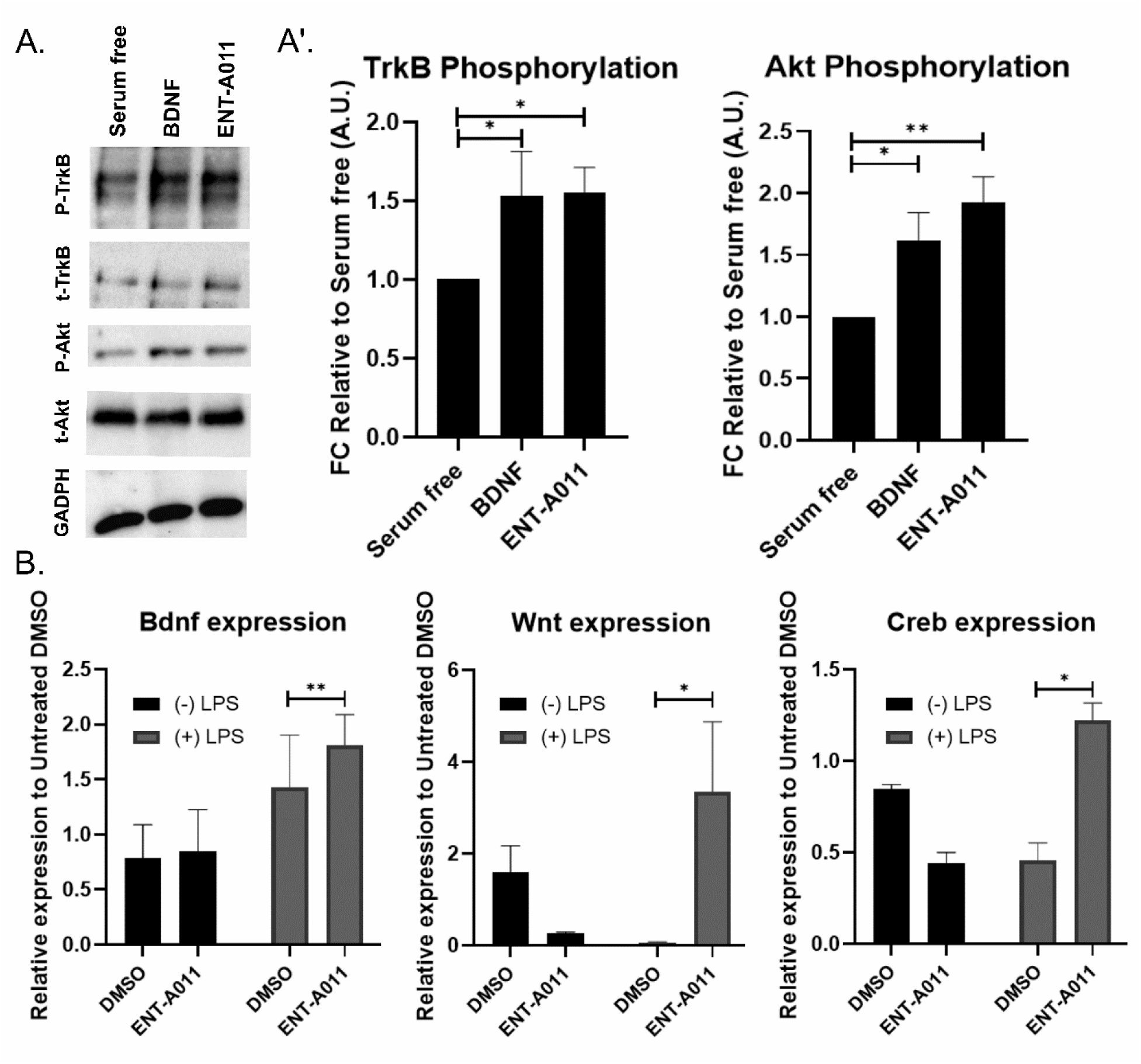
Effect of ENT-A011 on mouse astrocytes. The compound induces TrkB and its downstream target Akt phosphorylation after 20 minutes treatment in astrocytes. Representative blots (A) and quantification (A’) of 4 independent experiments are shown. Error bars represent S.E.M., Student’s t-test against Control; * p<0.05; *** p<0.001. B. ENT-A011 increases Bdnf, Wnt and Creb expression (detected by RT-qPCR) after LPS treatment in astrocytes. N=3-4, error bars represent S.E.M., Student’s t-test against Control; *<0.05, **<0.01.

### ENT-A011 promotes proliferation, survival and differentiation of adult primary hippocampal neural stem cells

Considering that the results presented above clearly demonstrated the TrkB-mediated effects of ENT-A011, we sought to investigate effects on adult neurogenesis. Thus, we used adult hippocampal neural stem cells isolated from mice and performed a set of complementary proliferation, survival and differentiation assays. First, we confirmed that ENT-A011 treatment leads to a significant increase in NSC proliferation rates (Figure 3A, A’) similar to BDNF (FC in proliferation rates equal to 1.92 compared to the untreated control for ENT-A011 and 1.97 for BDNF).

**Figure 3.**
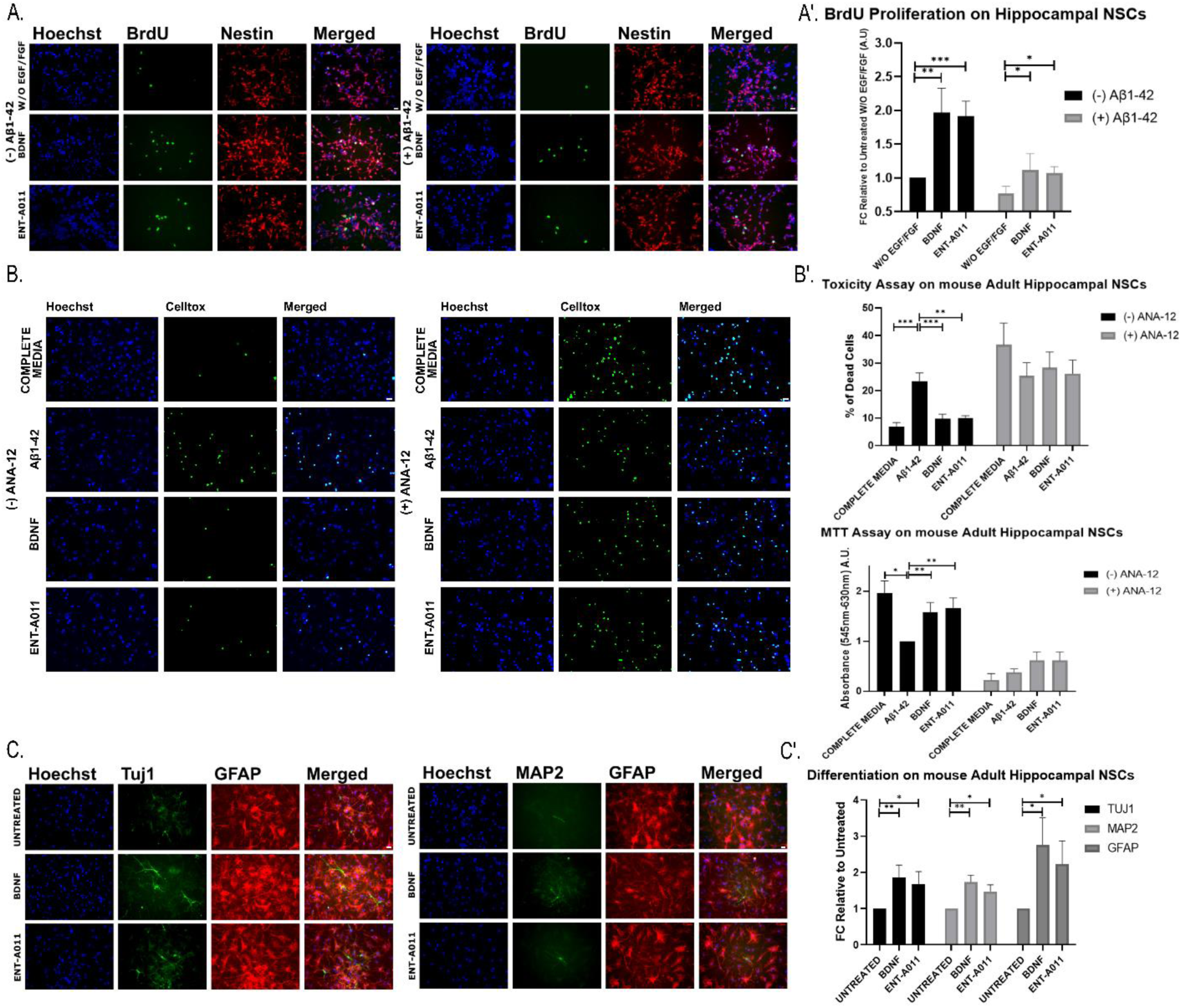
ENT-A011 induces neurogenesis on mouse adult hippocampal NSCs. A, A’. ENT-A011 increases proliferation in adult hippocampal NSCs. Representative fluorescence microscopy images of Immunostaining for BRDU and Nestin in adult hippocampal NSCs treated with BDNF or ENT-A011 for 24hrs with or without Aβ42 (A). Quantification of BRDU fluorescence change with or without Aβ42 (A’). Quantification represents BRDU positive cells normalized against Nestin positive cells. N=4, error bars represent S.E.M., Student’s t-test against Control; *<0.05, **<0.01, ***<0.001. B, B’. The compound rescues adult hippocampal NSCs from Aβ-induced cell death. Representative fluorescence microscopy images of Celltox cytotoxicity assay on hippocampal NSCs in the presence of Aβ42, treated with BDNF or ENT-A011, with or without the TrkB inhibitor ANA-12 for 24hrs (B). Quantification of fluorescence change and MTT levels after BDNF or compound treatment, with or without ANA-12 (B’). Quantification represents Celltox positive cells normalized against Hoechst positive cells. N=4, error bars represent S.E.M., Student’s t-test against Control; *<0.05, **<0.01, ***<0.001. C, C’. ENT-A011 promotes differentiation of adult hippocampal NSCs. Representative fluorescence microscopy images of Immunostaining for Tuj1 and GFAP in adult hippocampal neural stem cells treated with BDNF or ENT-A011 for 10 days (C). Fold change in expression of *TUJ1* (N=6 for untreated and BDNF, N=5 for ENTA-011), *MAP2* (N=5) and *GFAP* (N=4) assessed by RT-qPCR in BDNF or ENT-A011 treated adult hippocampal NSCs relative to untreated control cells. (C’). Scalebars = 20μm

We then assessed the effect of the compound on NSC survival after Aβ42 induced cytotoxicity. This system aims to mimic the conditions of the accumulation of oligomeric Aβ (with Aβ42 being the most toxic Aβ isoform), as a major described cause of neuronal cell death and reduced NSC proliferation in Alzheimer’s Disease (AD), especially in the hippocampal region which is the primary target of the disease. For this purpose, NSCs were treated with BDNF or ENT-A011 in the absence or presence of preformed Aβ42 oligomers (10μM). As shown in Figure 3A, A’, Aβ42 treatment led to a significant decrease in proliferation levels compared to the untreated control (0.77 FC relative to the untreated control in absence of Aβ42), while both BDNF and ENT-A011 were able to counter this toxic effect (FC 1.08 for ENT-A011 and 1.12 for BDNF relative to the untreated control in absence of Aβ42). In addition, we tested the ability of the compound to protect NSCs from cell death and restore reduced mitochondrial activity following Aβ42 toxicity. Both BDNF and ENT-A011 treatments successfully countered Aβ42-induced cell death, leading to apoptotic cell levels comparable to that in cells cultured in the absence of Aβ42 (23.44% of total cells were apoptotic in Aβ42, 9.82% for BDNF, 9.90% for ENT-A011, 6.91% for complete media without Aβ42) (Figure 3B, B’).

In parallel, the MTT assay was used to examine mitochondrial activity, showing higher absorbance and activity after treatment with ENT-A011 compared to the Aβ42 control, with a comparable increase to BDNF (FC compared to Aβ42 control 1.68 for ENT-A011 and 1.58 for BDNF). Furthermore, we performed both toxicity assays in the presence of the TrkB inhibitor, ANA-12. These experiments showed an abolishment of the effects of both BDNF and compound, supporting this neuroprotective action of the compound was also mediated through the TrkB receptor (Figure 3B, B’).

Finally, we tested the effect of the compound on NSC differentiation. NSCs were cultured for 10 days in the absence of proliferating factors EGF and FGF, then they were treated every two days with BDNF or ENT-A011, with the 10day protocol followed by immunostaining for TUJ1, MAP2 and GFAP markers for neuro-and glio-genesis. This showed that both BDNF and ENT-A011 promoted NSCs differentiation towards neurons and astrocytes (Figure 3C). Similarly, we observed an increase in mRNA levels by RT-PCR (QPCR) as depicted in Figure 3C’ (*Tuj1*: FC 1.67 for ENT-A011 and 1.86 for BDNF, *Map2*: FC 1.48 for ENT-A011 and 1.73 for BDNF, *Gfap*: FC 2.23 for ENT-A011 and 2.76 for BDNF).

### ENT-A011 promotes the proliferation and survival of embryonic primary cortical neural stem cells

Based on the results in adult hippocampal NSCs, we tested the effect of the compound on the proliferation of embryonic neural stem cells from the cortex. As shown in Figure 4A, A’, ENT-A011 drove a significant increase in proliferation of embryonic cortical NSCs (FC 1.34 compared to the untreated control), and counteracted the Aβ42-induced reduction of proliferation (increase from FC 0.72 to 1.01 after ENT-A011 treatment in presence of Aβ42 compared to the untreated control). Additionally, the compound reduced cell death levels similarly to BDNF (10.33% of total cells are apoptotic for ENT-A011 and 9.63% for BDNF compared to 22.6% in the untreated control) and promoted mitochondrial activity (FC in MTT absorbance of 2.24 for ENT-A011 and 2.18 for BDNF). Again, compound action of the compound in both assays was abolished in the presence of ANA-12, supporting it is mediated through TrkB signaling (Figure 4B, B’).

**Figure 4.**
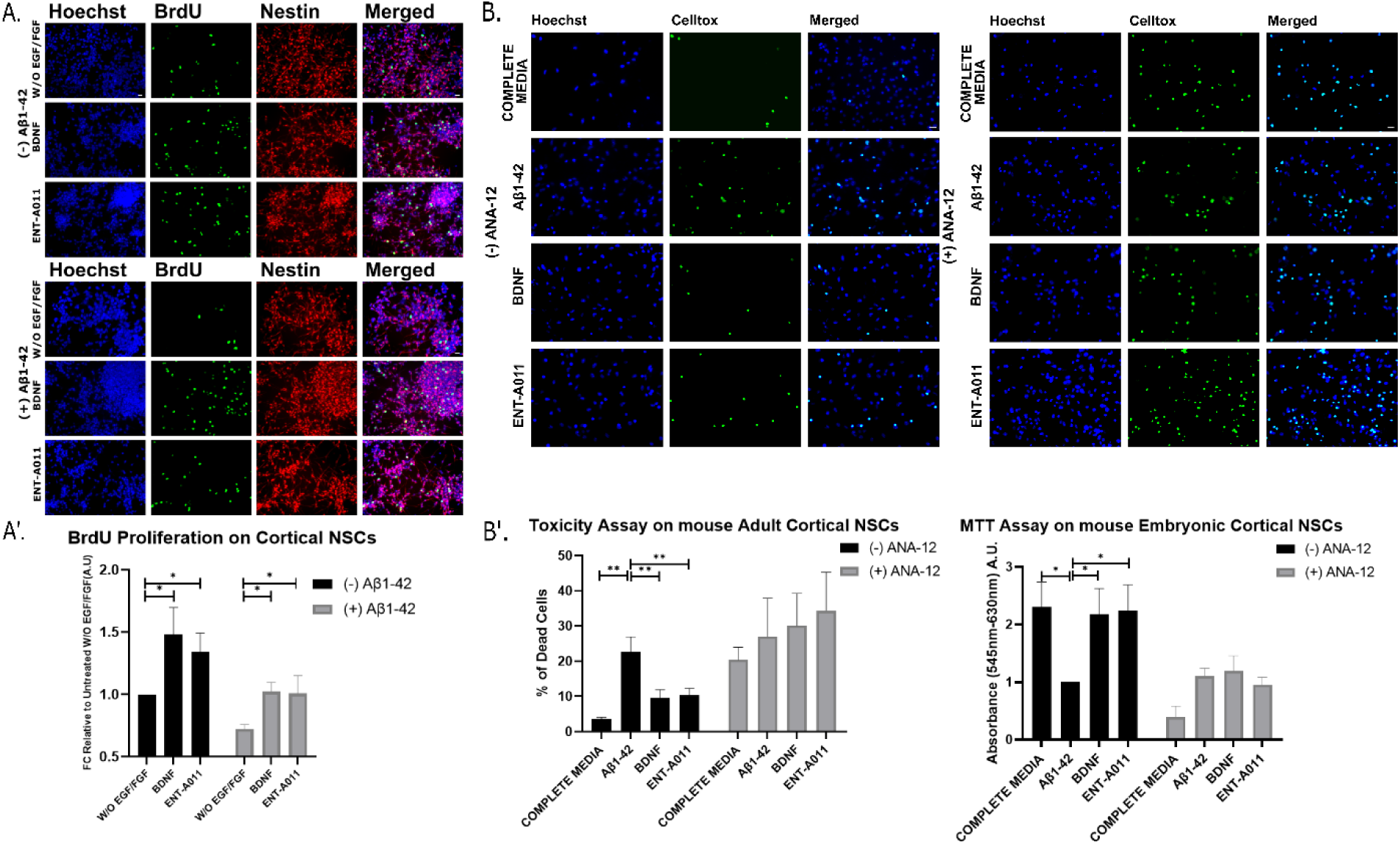
ENT-A011 has positive effect on mouse cortical NSCs. A, A’. ENT-A011 increases proliferation on cortical NSCs, Representative fluorescence microscopy images of Immunostaining for BRDU and Nestin in adult hippocampal NSCs treated with BDNF or ENT-A011 for 24hrs with or without Aβ42 (A). Quantification of BRDU fluorescence change with or without Aβ42 (A’). Quantification represents BRDU positive cells normalized against Nestin positive cells. N=4, error bars represent S.E.M., Student’s t-test against Control; *<0.05, **<0.01, ***<0.001. B, B’. The compound rescues cortical NSCs from Aβ-induced cell death. Representative fluorescence microscopy images of Celltox cytotoxicity assay on hippocampal NSCs in the presence of Aβ42, treated with BDNF or ENT-A011, with or without the TrkB inhibitor ANA-12 for 24hrs (B). Quantification of fluorescence change and MTT levels after BDNF or compound treatment, with or without ANA-12 (B’). Quantification represents Celltox positive cells normalized against Hoechst positive cells. N=4, error bars represent S.E.M., Student’s t-test against Control; *<0.05, **<0.01, ***<0.001. C, C’. Scalebars = 20μm

### ENT-A011 protects hippocampal neurons from Aβ42 cytotoxicity and promotes neurite outgrowth of cortical neurons

Complementary to the anti-amyloid activity of ENT-A011 in hippocampal neural stem cells, we were also interested to test the action of the compound in counteracting the Aβ42-induced hippocampal neuron death and synapse loss. Oligomeric Aβ reduced neuronal trophic support, which is partly caused by neurotrophic signaling deregulation. It stands to reason that a small molecule that selectively activates TrkB signaling and promotes neuronal survival could mitigate the negative effects of toxic Aβ, particularly in the early stages of the disease, acting against cell death and synapse degeneration. To assess the anti-amyloid neuroprotective effects of ENT-A011, we cultured neurons isolated from the hippocampus of E17.5 mouse embryos, a brain area that is severely impacted in Alzheimer’s disease. Primary neurons were cultured for 16-18 DIV before being treated for 48 hours with oligomeric Aβ and ENT-A011. After cell fixation, apoptosis was determined using the TUNEL assay. ENT-A011-treated cells showed significantly lower apoptosis (20.5% ± 4.1%) than Aβ-treated cells (41.5% ± 4.2%), at comparable levels to control cells (18.7% ± 2.8%) (Figure 5A, A’). We then tested the efficacy of ENT-A011 to protect against Aβ-induced synapse degeneration, by treating primary hippocampal neurons for 4 hrs with Aβ and ENT-A011. As seen in Figure 5B, B’ synaptophysin immunostaining depicted that the ENT-A011-treated group shows a comparable number of synapses to the control group (FC 1.05 ±0.06 of control), while the Aβ-treated group had significantly reduced synapses (FC 0.78 ± 0.03 of control). These findings suggest that ENT-A011 protects primary hippocampal neurons from oligomeric Aβ-induced cell death and against synapse degeneration.

**Figure 5.**
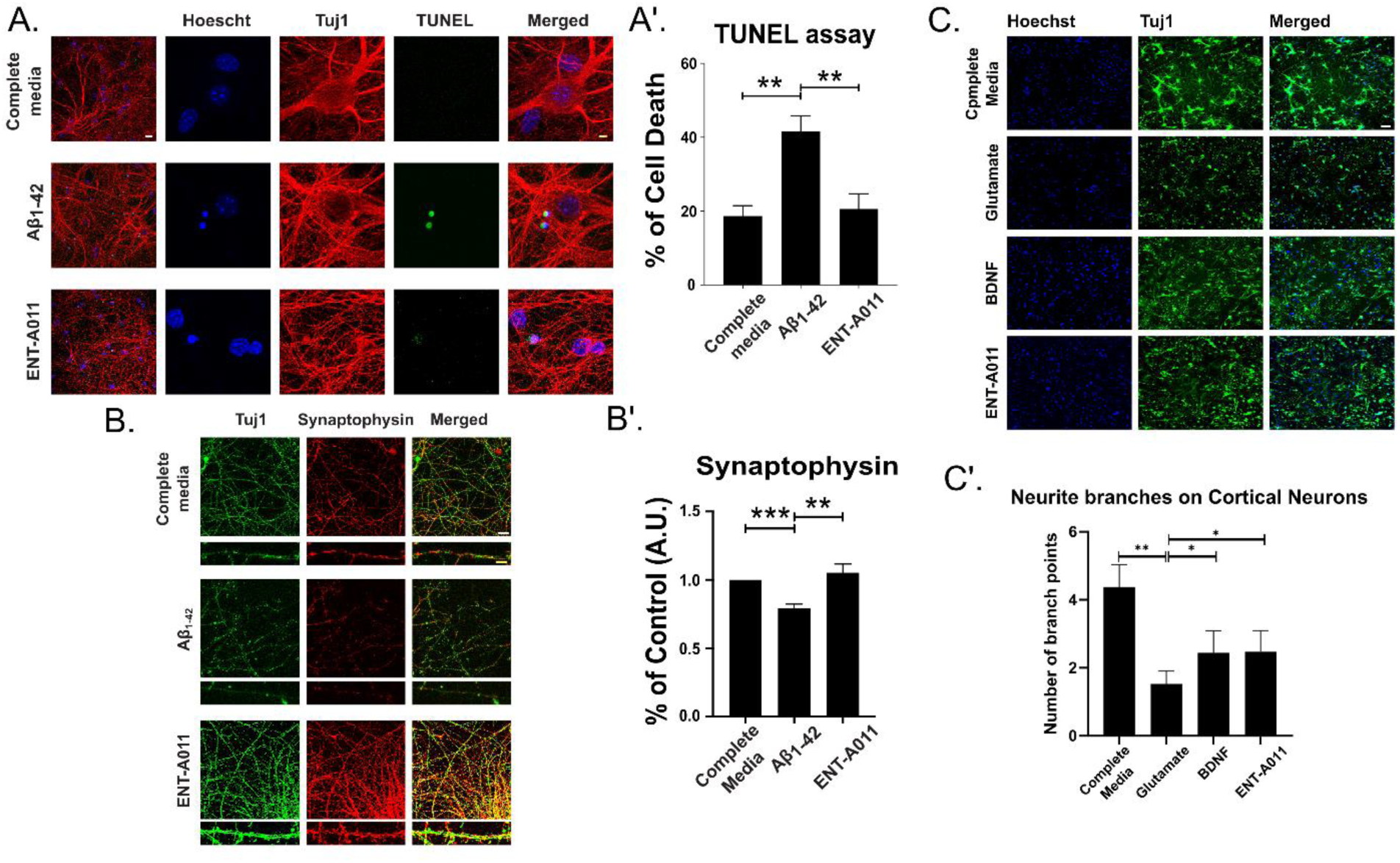
ENT-A011 protects mouse hippocampal neurons and rat cortical neurons against Aβ42 and glutamate toxicity respectively. A, A‘, ENT-A011 reduces Aβ42 induced hippocampal neuron cell death. Primary hippocampal neurons were treated with Aβ42 for 48 hours in presence of ENT-A011 and cell death was quantified using TUNEL assay. Representative images from 4 independent experiments. Data are shown as SEM. One-way ANOVA, Tukey’s Test correction: ** p < 0.01; *** p < 0.001. B, B’, ENT-A011 protects against Aβ42-induced hippocampal synapse loss. Primary hippocampal neurons were treated with Aβ42 for 4 hours in presence of ENT-A011, followed by staining against synaptophysin to assess synapse number. Total Synaptophysin area was normalized to total Tuj1 area in each image. Representative images from 4 independent experiments. Data are shown as SEM. One-way ANOVA, Tukey’s Test correction: ** p < 0.01. ENT-A011 reverts the decrease of neurite branches caused by glutamate on rat Cortical neurons. Representative fluorescence microscopy images of Immunostaining for Tuj1 in cortical neurons treated with BDNF or ENT-A011 for 1 hour before glutamate treatment (C). Quantification of neurite branches (C’). N=3, error bars represent S.E.M., Student’s t-test against Control; *<0.05, **<0.01, ***<0.001. White Scalebars = 20μm, Yellow Scalebars = 5μm

In concert with Aβ42 toxicity, excitotoxicity caused by high concentrations of glutamate and abnormal activation of glutamate receptors (NMDA) is also known to be an important event in the neuropathology of AD, and a therapeutic target of the drug Memantine, a NMDA-antagonist. Thus, to investigate the effect of ENT-A011 upon glutamate treatment, we cultured primary rat cortical neurons and tested how neurite branches were affected after treatment. Both BDNF and ENT-A011 promoted neurite outgrowth in cortical neurons subjected to a glutamate excitotoxicity condition (Figure 5C, C’), calculated as a change in neurite branches per neuron (average of 2.44 branch points for BDNF and 2.47 for ENT-A011 compared to 1.53 for the control treated only with glutamate). Taken together, results in hippocampal and cortical cell populations, both of mature and stem cell origin, support the hypothesis the compound may have significant therapeutic potential against major pathological conditions of AD.

### ENT-A011 increases the proliferation and survival from Aβ42-induced cell death of human neural progenitor cells, derived from healthy and AD-induced pluripotent stem cell lines

The ensemble of functional assays in various TrkB-expressing cell lines and different neural stem cell populations from mice supported the hypothesis that ENT-A011 has TrkB receptor mediated activity comparable to that of BDNF in both healthy and conditions simulating AD. Nevertheless, mouse models may differ importantly from homologous human cell populations and systems. Thus, we designed a human cell based experimental setup, as a platform with greater human translational potential for applications of a BDNF mimetic for AD research. In this setup, we used three different human iPSC lines, two from healthy individuals and one from an APOE4-AD individual. We carried out differentiation of these lines towards cortical neuronal progenitor cells (NPCs), using an established protocol, that mimics corticogenesis *in vitro*^79^. Treatment of human NPCs at differentiation days 25-27 with ENT-A011 for 48 hours was able to drive an increase in proliferation as seen in Figure 6A, A’ (72.23% of total cells were BRDU after ENT-A011 treatment compared to 65.02% in untreated control for the first healthy line, 70.95% in ENT-A011 compared to 61.44% in the control for the second healthy line and 81.47% in ENT-A011 compared to 72.01% in the control for the APOE4 line). Furthermore, NPCs from the three different lines at the same differentiation days were cultured and subjected to Aβ42 toxicity, followed by compound treatments. ENT-A011 treatment was able to counter Aβ42 induced toxicity in human NPCs, leading to a reduction in Celltox assay cell death levels and an increase in MTT assay mitochondrial activity, similarly to BDNF (Figure 6B, B’).

**Figure 6.**
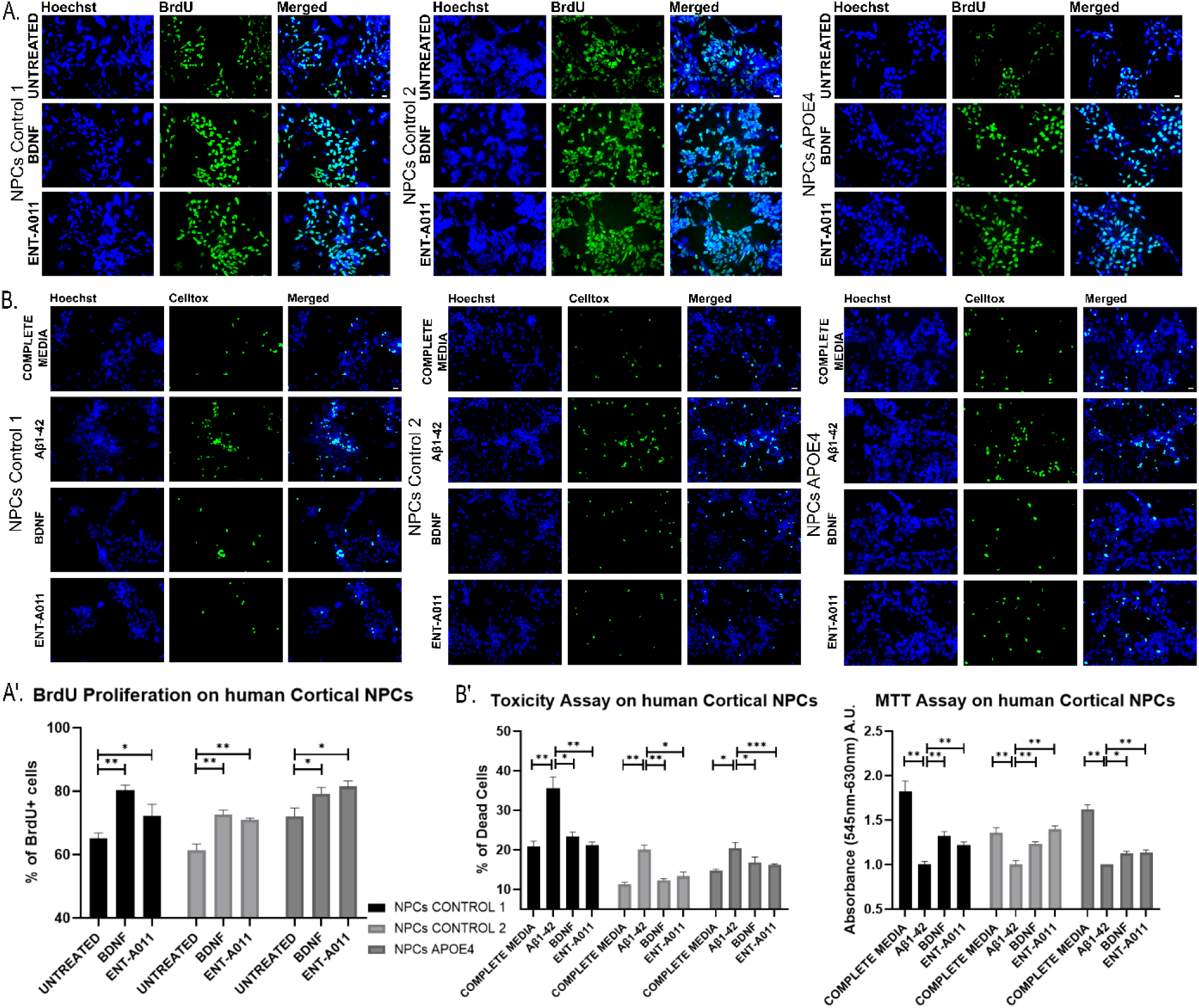
ENT-A011 effect on human cortical Neural Progenitor Cells (NPCs). Change in BRDU+ cells (calculated as percentage of total Hoechst+ cells) after BDNF or ENT-A011 treatment for 48hrs in NPC derived from 3 human induced pluripotent stem cell (iPSC) lines. Representative fluorescence microscopy images of Immunostaining for BRDU and Hoechst (A) and quantification of BRDU fluorescence change (A’). B, B’. NPC derived from 3 human iPSC lines were treated with Aβ1-42 oligomers and BDNF or ENT-A011 for 48hrs and compound effect on reducing toxicity was assessed via the Celltox cytotoxicity assay. Representative fluorescence microscopy images of Celltox cytotoxicity assay in the presence of Aβ42, treated with BDNF or ENT-A011 for 24hrs (B). Quantification of fluorescence change and MTT levels after BDNF or compound treatment, with or without ANA-12 (B’). Quantification represents Celltox positive cells normalized against Hoechst positive cells. Three different lines were differentiated, N=3, error bars represent S.E.M., Student’s t-test against Control; *<0.05, **<0.01, ***<0.001. Scalebars = 20μm

Finally, we wanted to obtain a quantitative and global assessment of the gene networks underlying ENT-A011 action and if they are comparable to BDNF. For this purpose, we carried out 3’ quant mRNA sequencing from total mRNA extracted from human APOE4 iPSC derived NPCs (day 27) treated with BDNF, ENT-A011 or from untreated controls. Differential expression analysis (DEA) using the EDGER package revealed a core network of 140 genes that were differentially regulated in the same way by both ENT-A011 and BDNF, amounting to nearly 60% of all genes differentially regulated by BDNF (233 genes, Fig. 7a). Unexpectedly, ENT-A011 treatment led to differential expression of 7.9-fold more target genes (1840) than BDNF (Figure 7b). Focusing on the commonly differentially regulated genes, we note a very reproducible pattern in expression changes after ENT-A011 and BDNF (Figure 7C) and we confirmed a significant positive correlation between the effect of BDNF and that of ENT-A011 on the expression of common target genes, as seen in Figure 7D (Spearman Correlation coefficient 0.9, pvalue 2.2e^-16^). Genes upregulated by either treatment were also found to be associated with shared enriched ontology terms, including “neurotransmitter secretion”, “synapse organization” and “regulation of cell growth” (Figure 7e). Genes upregulated by ENT-A011 exclusively are associated with additional processes related to neuronal development and function such as “cerebral cortex development”, “regulation of neurogenesis” and “signaling by NTRKs” (Figure 7e).

**Figure 7.**
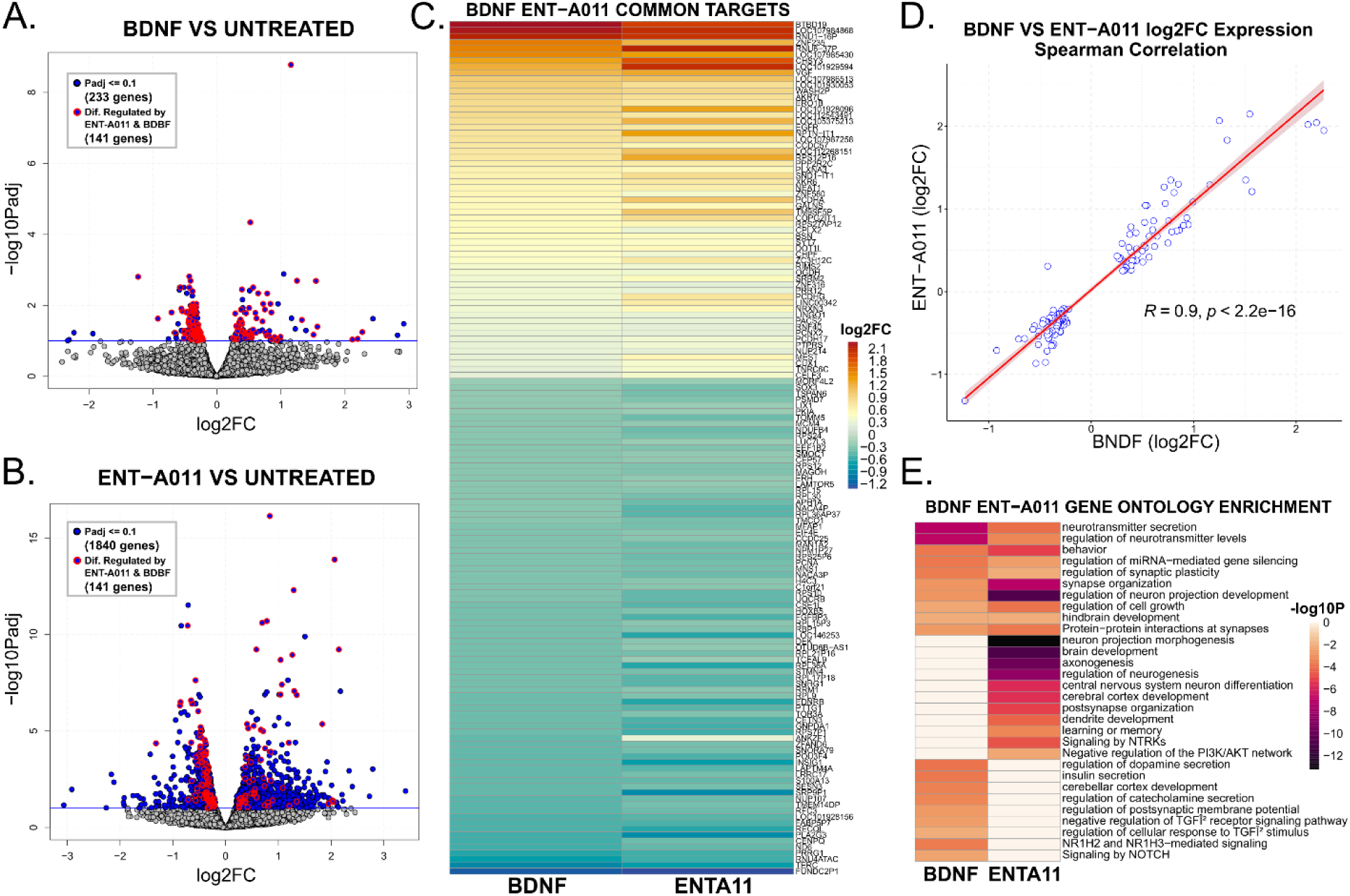
Differential expression analysis of ENT-A011 and BDNF treated APOE4 human Cortical Neural Progenitor Cells (NSCs) compared to untreated control cells. a. Volcano plot of differentially regulated genes after BDNF treatment compared to untreated control cells. Negative log10 adjusted Pvalues (y axis) are plotted against log2 fold change in expression for each gene. Genes with Padj lower or equal to 0.1 (blue horizontal line) are highlighted in bold blue colour and genes that are also differentially regulated by ENT-A011 (Padj<=0.1) are highlighted with red circles. **b.** Volcano plot of differentially regulated genes after ENT-A011 treatment compared to untreated control cells. Negative log10 adjusted Pvalues (y axis) are plotted against log2 fold change in expression for each gene. Genes with Padj lower or equal to 0.1 (blue horizontal line) are highlighted in bold blue colour and genes that are also differentially regulated by ENT-A011 (Padj<=0.1) are highlighted with red circles. **c.** Heatmap of log2 Fold Change in expression for genes differentially regulated by both BDNF and ENT-A011, ranked by decreasing log2FC values in BDNF treated cells. **d.** Scatter plots of log2 fold change in gene expression after BDNF (x axis) or ENT-A011 (y axis) treatment. Spearman correlation coefficient (0.9) and pvalue (2.2e^-16^) are displayed next to line of best fit. **e.** Heatmap of negative log10 Pvalues for gene ontology enrichment analysis of upregulated genes (Padj <=0.1) after BDNF (left column) or ENT-A011 treatment. Unique terms to either treatment group are plotted in pale orange in the other group.

## DISCUSSION

BDNF is the most widely distributed neurotrophic growth factor in the central nervous system (CNS), with crucial roles for CNS growth and development. BDNF has been shown to promote neurogenesis, both in the embryonic and adult brain, and is essential for neural stem cell proliferation, differentiation and survival^21, 24, 92, 93^. In recent years, compelling evidence has suggested neurogenesis impairment in Alzheimer’s disease to be linked to decreased levels of the BDNF in the hippocampus and the cortex^29^. Consequently, various studies have investigated the therapeutic potency of BDNF in neurodegenerative diseases and especially in AD^21, 32^. However, its inability to penetrate the blood-brain barrier, minimal diffusion in tissues and sensitivity to proteolysis hinder the potential of BDNF as a drug candidate for AD. Aiming to overcome these shortcomings and based on our previous studies on steroidal neurotrophin mimetics, we designed and synthesized compound ENT-A011 which shared the beneficial functions of BDNF in TrkB dependent cell lines and neural cell populations, but not its limitations. In the present study, we characterize ENT-A011 ability to rescue TrkB-expressing neuronal cells, an effect comparable to that of BDNF. Importantly, our experiments show that this novel compound has neurogenic and neuroprotective properties, not only in mouse neuronal cell populations, but most importantly in human neural progenitor cells derived from healthy and AD donors.

ENT-A011 induces phosphorylation of TrkB, but not TrkA or TrkC receptors while it does not activate the pan-neurotrophin p75^NTR^ receptor pathway (Figure S8). Molecular modeling suggested two possible binding sites for ENT-A011 at the interface of the extracellular TrkB domain with the neurotrophin. ENT-A011 also activates TrkB receptor downstream signaling targets, increasing Akt phosphorylation, a kinase pathway associated with cell survival. Indeed, the compound reduces cell death levels after serum deprivation in survival assays in NIH-3T3 cells, stably transfected with TrkB, an effect abolished after treatment with TrkB inhibitor ANA-12, providing further evidence that the compound specifically protects cells through TrkB receptor activation. In this first round of screening in TrkB-expressing cell lines, we also show dose-dependent action of the compound by using a nomad biosensor cell system produced by Innoprot. Having confirmed that ENT-A011 can activate the TrkB receptor and the downstream pathway and its functional role in survival, we then investigated the actions of the compound on mouse primary astrocytes. These cells naturally express TrkB expressing population, playing an important role in neurogenesis. The interaction of astrocytes with neural stem cells supports proliferation, differentiation and synapse formation of the latter, acting as a major source of BDNF^23, 94–98^. ENT-A011 showed the ability to induce phosphorylation of the full length TrkB receptor isoform in primary astrocytes, and it was also able to induce *Creb, Wnt* and *Bdnf* expression after LPS-induced astrocyte activation^99^. LPS leads to reduction in cell proliferation and the production of new neurons^100, 101^. It is of note that the levels of hippocampal BDNF and phospho-TrkB were shown to decrease after astrocyte activation in LPS treated mice^102^.

Activation of *Creb* in astrocytes is involved in proliferation, survival, maturation and development of neural stem and progenitor cells and correlates with the expression and activation of BDNF^103, 104^. In addition, the *Wnt* family is expressed by astrocytes and Wnt receptors are present in neural stem cells. Wnt signaling has been shown to hold roles in enhancing proliferation, the induction of neuronal differentiation, fate-commitment, development and maturation of neural stem cells, progenitor cells and newborn neurons^94, 105, 106^. It is also worth noticing that astrocytes express both the TrkB full length receptor and the TrkB truncated form of the receptor depending upon their maturation stage, with both types of the receptor been activated by BDNF, either by phosphorylation of the tyrosine residue for the full length isoform or through calcium release for the truncated form^107–109^. Therefore, our results confirm the ability of ENT-A011 to activate the TrkB pathway in a naturally expressing TrkB cell population and provide evidence on its effect on the expression of genes that are associated with astrocyte neurogenic and its neuroprotective action.

Alzheimer’s disease is characterized by the accumulation of toxic oligomeric amyloid-β and neuronal loss in hippocampus and cortex. Oligomeric Amyloid-β, especially Aβ42, reduces neuronal proliferation and differentiation, an event associated with reduced hippocampal neurogenesis. Additionally, Aβ42 induces apoptosis at a concentration of 10μM in neural progenitor cells^110, 111^. Aβ affects neurogenesis in mouse and human models of Alzheimer’s disease^112–114^. An increasing number of studies suggest that induction of adult neurogenesis before the progression of AD may be a potential therapeutic or prevention strategy earlier in life^115, 116^. We thus examined the effect of ENT-A011 on the proliferation and survival of mouse hippocampal and cortical neural stem cells, in the absence or the presence of Aβ42, compared to the activity of BDNF. ENT-A011 was able to significantly reduce levels of cell death in survival assays and countered the decline of cell proliferation after Aβ42 treatment. Moreover, ENT-A011 induced hippocampal neural stem cell differentiation towards neurons and astrocytes. These findings considered together, strongly support the ability of the compound to promote neurogenesis mimicking BDNF. It is well established that Amyloid-β also impairs NSC viability by disrupting mitochondrial function^117^. Interestingly, ENT-A011 treatment reversed the reduction in metabolic activity caused by Aβ42 treatment, in a manner comparable to BDNF.

Access to human induced pluripotent stem cells in latest years has offered the opportunity to study neurogenesis in a human system. Such new translational platforms for human disease, open a unique channel for studying the effects of BDNF on human neural stem cells and neurons and provide a system to screen potential neurogenic drugs for neurodegenerative diseases and especially AD. Previous work has shown that BDNF induces the proliferation of human neural progenitor cells from both control and AD iPSC lines ^118^. In the present study, we show that ENT-A011, as a synthetic BDNF mimetic, has the ability to promote proliferation and mitochondrial activity, as well as decrease Aβ42 induced cell death in neural progenitor cells derived from iPSCs from healthy controls and an APOE4 cell line. Interestingly, BDNF was found decreased in human APO4 carriers^119^, and hippocampal progenitor cell proliferation was decreased in mice with the human APOE4^120, 121^, while NPCs from APOE4 iPSC-derived neurons showed impaired capacity for proliferation^122^. We thus compared the effects of BDNF and ENT-A011 treatment on human NPCs derived from iPSCs from the APOE4 line, comparing their actions on target gene network activation, using RNA sequencing. Differential expression analysis showed that ENT-A011 activates a downstream gene network largely overlapping with the downstream network activated by BDNF. Indeed, nearly 60% gene targets were upregulated and downregulated by the compound in a similar fashion as BDNF. Upregulated targets after ENT-A011 treatment are associated with regulation of cell growth, axonogenesis, regulation of neurogenesis and signaling by NTRKs, providing valuable support of the action of ENT-A011 in human neuronal progenitors.

Genes co-upregulated by BDNF and ENT-A011 include various targets of interest associated with neural development and function, as well as neuronal disorders and AD pathology. Complexin 2 (*CPX-2*) modulates activity-induced BDNF release in hippocampal neurons^123^, while cut like homeobox 1 (*CUX1*) regulates spine development, dendritic branching and synaptogenesis in cortical neurons^124^. Reduced neurexin 3 (*NRXN3*) expression has been associated with increased inflammation in AD and Aβ oligomer interaction with neurexins leading to inhibition of presynaptic differentiation^125, 126^. Downregulation of Protein Tyrosine Phosphatase Receptor Type S (*PTPRS*) results in increased TrkB phosphorylation and long-term memory impairments in mice^127^. A co-upregulated target of particular interest is presynaptic organizer bassoon (*BSN*), as not only do BSN mutant mice have increased BDNF expression, but this leads to hippocampal enlargement because of increased proliferation and reduced apoptosis of new neurons in the dentate gyrus^128^. Finally, human haploinsufficiency of Proline Rich 12 (*PRR12*) has also been linked to neurodevelopmental abnormalities^129^. Additionally, specific upregulated targets of ENT-A011 provide further genes of interest, including t-Box Brain Transcription Factor 1 (*TBR1*), a regulator of neural stem cell differentiation that is expressed during cortical development and in the adult hippocampus^130, 131^. Moreover, DMRT Like Family A2 (*DMRTA2*) regulates NPC maintenance during cortical development and is necessary for early cortical neurogenesis in mice^132, 133^. Another member of the Protein Tyrosine Phosphatase Receptor family, *PTPRD*, upregulated by ENT-A011 is a regulator of developmental neurogenesis through TrkB^134^. RAR Related Orphan Receptor A (*RORA*) has been identified as a BDNF interaction partner and increased RORA expression has been shown in the AD hippocampus^135^. A striking target of ENT-A011 is also amyloid Beta Precursor Like Protein 1 (*APLP1*), a member of the amyloid precursor protein family involved in brain development that is directly cleaved by γ-secretase, but whose derivative ALP-1 peptides are not toxic to neuronal cells, unlike APP derived Aβ^136, 137^. Finally, ENT-A011 upregulated targets also include BDNF/TrkB signalling members, such as docking protein 5 (*DOK5*) that act as a substrate of *TrkB*^138^, NCK Adaptor Protein 2 (*NCK2),* whose interaction with TrkB has been shown to be promoted by BDNF in cortical neurons^139^ and BCL11 Transcription Factor B (*BCL11B*) that acts as an upstream regulator of multiple other BDNF signaling genes^140^.

In this study we performed a broad characterization of the ability of a new candidate therapeutic to promote neurogenesis and neuroprotection using a complement of adult and embryonic stem cell models. By combining relevant mouse cell lines with human iPSC derived NPCs, we strived to create a robust in-vitro testing platform with translational value for neurogenic drug development, in line with promoting the 3R principles of replacing, reducing and refining animal use in research. This work presents a novel TrkB agonist, ENT-A011, that exhibits high neurogenic and neuroprotective action on par with the endogenous ligand BDNF, while having advantageous drug like properties for putative therapeutic applications that includes its small size, a lipophilic nature and appropriate metabolic profile.

## Aknowledgments

RNA-seq library preparation, sequencing and preliminary analyses were performed at the Genomics Facility of IMBB FORTH, Heraklion, Crete, Greece. We would like to thank Isbaal Ramos, Patricia Villace and Rosa M. Mella for hosting arrangements and technical assistance with experiments at Innoprot, Bilbao. We would also like to thank Paula Rocktaeschel and Emma M. Hayes for support with iPSC experiments in Oxford and coordinating shipping arrangements. Finally, we would like to thank Canelif Yilmaz for gene expression experiments in Dresden and Alexandros Tsimpolis for assistance and guidance on astrocyte culture and RT-qPCR optimization.

