## Supplementary Data for "Comprehensive characterization of the neurogenic and neuroprotective action of a novel TrkB agonist using mouse and human stem cell models of Alzheimer’s Disease"

Synthesis of ENT-A011 and ENT-A012

*Synthesis of (E)-(3β-Hydroxy-5-androsten-17-ylidene) ethyl ester (1)*

To a solution of DHEA (2.0 g, 6.94 mmol) and triethyl phosphonoacetate (15.46 mL, 77.9 mmol) in anhydrous tetrahydrofuran (THF)/absolute ethanol 1:1 (21.4/21.4 mL) was added dropwise at 25 °C a solution of sodium ethoxide (EtONa) in absolute ethanol (prepared from 1.59 g Na in 30 mL ethanol). The resulting mixture was refluxed overnight. Subsequently, the reaction was cooled to 0 °C and was carefully quenched with water and acidified using 10% aq. HCl until completion of formation of a precipitate. The solid was filtered under reduced pressure, washed with water (30 mL × 3 times) and petroleum ether 40–60 °C (30 mL × 3 times), and air-dried. The solid was dissolved in $CH_2Cl_2$ and the solution was dried over anhydrous $Na_2SO_4$ and filtered. The filtrate was then evaporated under reduced pressure to afford compound **1** (2.39 g, 96% yield) as a pure white crystalline solid. The compound was used in the next step without further purification. Mp: 178–180 °C; $[\alpha]_D^{25}$= −78° (c = 0.006 g/mL, $CHCl_3$); $R_f$: 0.5 (petroleum ether 40–60 °C/acetone 80:20); $^1$H NMR (600 MHz, $CDCl_3$): $\delta$ 0.84 (s, 3H), 1.03 (s, 3H), 1.28 (t, $J$ = 7.1 Hz, 3H, $OCH_2$*$CH_3$*), 1.31–2.34 (m, 18H), 2.78–2.90 (m, 2H), 3.48–3.57 (m, 1H, 3$\alpha$-H), 4.15 (q, $J$ = 7.1 Hz, 2H, O*$CH_2$*$CH_3$), 5.36 (d, $J$ = 5.0 Hz, 1H, 6-H), 5.55 (bs, 1H, 20-H); $^{13}$C NMR (150 MHz, $CDCl_3$): $\delta$ 14.5, 18.4, 19.6, 21.1, 24.6, 30.6, 31.7, 31.8, 35.3, 36.8, 37.4, 42.4, 46.2, 50.4, 54.0, 59.7, 71.7, 71.8, 108.8, 121.5, 140.9, 167.6, 176.3; APCI-HRMS: m/z calculated for $[M+H]^+$ $C_{23}H_{35}O_3$ 359.2581, found 359.2579.

*Synthesis of (E)-[3β-(t-butyldimethylsilyloxy)-5-androsten-17-ylidene] ethyl ester (2)*

To a solution of compound 1 (2.25 g, 6.28 mmol) in anhydrous THF (20 mL) imidazole (1.33 g, 19.5 mmol) and iodine (4.76 g, 37.5 mmol) were added at 0 °C and the mixture was stirred at 0 °C for 30 min. Subsequently, *tert*-butyldimethylsilyl chloride (1.05 g, 6.67 mmol) was added and the resulting mixture was stirred at 25 °C overnight. After completion of the reaction, the solvent was evaporated in vacuo and the residue was diluted and extracted using ethyl acetate (EtOAc). The organic layer was washed using sat. aq. $Na_2S_2O_4$, brine and dried over anhydrous $Na_2SO_4$. The solvent was removed *in vacuo* and the residue was purified by FCC (elution solvent: petroleum ether 40–60 °C/ethyl acetate 95:5) to afford compound 2 as a white solid (2.36 g, 93% yield). Mp: 100–102 °C; $[\alpha]_D^{25}$= −50° (c= 0.0056 g/mL, $CHCl_3$); $R_f$: 0.86 (petroleum ether 40–60 °C/EtOAc 80:20); $^1$H NMR (600 MHz, $CDCl_3$): $\delta$ 0.05 (s, 6H, $Si(CH_3)_2$), 0.83 (s, 3H), 0.89 (s, 9H, $C(CH_3)_3$), 1.02 (s, 3H), 1.28 (t, $J$ = 7.1 Hz, 3H, $OCH_2$*$CH_3$*), 1.32–2.31 (m, 17H), 2.79–2.89 (m, 2H) 3.42–3.54 (m, 1H, 3$\alpha$-H), 4.15 (q, $J$ = 7.1 Hz, 2H, O*$CH_2$*$CH_3$), 5.33 (d, $J$ = 5.3 Hz, 1H, 6-H), 5.54 (t, $J$ = 2.3 Hz, 1H, 20-H); $^{13}$C NMR (75 MHz, $CDCl_3$): $\delta$ -4.3, 14.5, 18.4, 19.6, 21.1, 24.6, 26.1, 30.6, 31.7, 31.8, 32.2, 35.4,

36.8, 37.5, 42.9, 46.2, 50.5, 54.0, 59.7, 72.7, 108.8, 120.9, 141.8, 167.6, 176.4; APCI-HRMS: m/z calculated for $C_{29}H_{49}O_3Si$ [M+H]$^+$ 473.3445, found 473.3443.

*Synthesis of (E)-3β-(t-butyldimethylsilyloxy)-pregna-5,17(20)-dien-21-ol (3)*

To a solution of compound **2** (2.20 g, 4.65 mmol) in anhydrous THF (94 mL) was added dropwise at −78 °C diisobutylaluminum hydride (DIBAL-H) [(1.0 M in hexane), 18.7 mL, 18.7 mmol]. The reaction mixture was stirred at −78 °C for 2.5 h and at 25 °C for an additional 1 h. After completion of the reaction, saturated aqueous $NH_4Cl$ (40 mL) was added at 0 °C and the solvent was evaporated in vacuo. EtOAc (30 mL) was added to the residue and the organic layer was washed using 10% aq. HCl (20 mL) and brine (40 mL), dried over anhydrous $Na_2SO_4$ and filtered. The filtrate was then evaporated under reduced pressure to afford compound 3 as a white pure crystalline solid (2.0 g, quantitative yield). Compound **3** was used in the next step without further purification. Mp: 129–131 °C; $[\alpha]_D^{25}$= −39° (c = 0.0054 g/mL, $CHCl_3$); $R_f$: 0.46, (petroleum ether 40–60 °C/EtOAc 80:20); $^1H$ NMR (600 MHz, $CDCl_3$): $\delta$ 0.06 (s, 6H, $Si(CH_3)_2$), 0.78 (s, 3H), 0.89 (s, 9H, $C(CH_3)_3$), 1.02 (s, 3H), 1.19–2.43 (m, 18H), 3.43–3.53 (m, 1H, 3$\alpha$-H), 4.09 (dd, $J$ = 12.2, 6.7 Hz, 1H, 21-H), 4.15 (dd, $J$ = 12.2, 6.7 Hz, 1H, 21-H) 5.25 (t, $J$ = 6.7 Hz, 1H, 20-H), 5.32 (d, $J$ = 5.1 Hz, 1H, 6-H); $^{13}C$ NMR (75 MHz, $CDCl_3$): $\delta$ −4.4, 14.3, 18.4, 18.7, 19.6, 21.1, 21.2, 24.5, 26.1, 26.3, 31.8, 31.9, 32.2, 35.8, 36.9, 37.5, 42.9, 44.0, 50.7, 72.7, 115.7, 121.0, 141.8, 155.9; APCI-HRMS: m/z calculated for $C_{21}H_{31}O$ [(M-TBS-$H_2O$)+H]$^+$ 299.2369, found 299.2367.

*Synthesis of (17S,20S)-3β-(t-butyldimethylsilyloxy)-17,20-methan-5-pregnene-21-ol (4)*

To a solution of compound **3** (1.8 g, 4.17 mmol) in anhydrous toluene (33.6 mL) was added at 25 °C diiodomethane ($CH_2I_2$) (1.8 mL, 21.6 mmol). The reaction mixture was then cooled to −78 °C and a solution of diethylzinc (0.9 M in hexane, 24 mL, 21.6 mmol) was added. The reaction mixture was then stirred at 25 °C for 2 h. After completion of the reaction (monitored by $^1H$ NMR), the reaction mixture was quenched using 10% aqueous HCl at 0 °C until pH 5.5, followed by extraction using EtOAc (30 mL × 3 times). The organic layer was washed with brine, dried over anhydrous $Na_2SO_4$ and the solvent was removed *in vacuo.* The residue was purified by FCC (petroleum ether 40–60 °C/EtOAc 90/10 → 85/15) to obtain compound **4** (1.1 g, 60% yield). Mp: 154–157° C; $[\alpha]_D^{25}$= −50° (c= 0.0036 g/mL, $CHCl_3$); $R_f$: 0.46 (petroleum ether 40–60 °C/EtOAc 80:20); $^1H$ NMR (600 MHz, $CDCl_3$): $\delta$ 0.06 (s, 6H, $Si(CH_3)_2$), 0.78 (s, 3H), 0.89 (s, 9H, $C(CH_3)_3$), 0.90–0.97 (m, 1H) 1.01 (s, 3H), 1.03–2.32 (m, 20H), 3.45–3.54 (m, 2H), 3.58–3.66 (m, 1H, 22-H), 5.33 (d, $J$ = 5.3 Hz, 1H, 6-H); $^{13}C$ NMR (75 MHz, $CDCl_3$): $\delta$ −4.4, 16.3, 17.1, 18.4, 19.6, 19.8, 20.6, 25.2, 26.1, 29.1, 32.1, 32.2, 32.7, 33.3, 36.3, 36.8, 37.5, 41.1, 42.9, 50.6, 54.9, 65.3, 72.7, 121.1, 141.7; APCI-HRMS: m/z calculated for $C_{28}H_{47}OSi$ [(M-$H_2O$)+H]$^+$ 427.3391, found 427.3386.

*Synthesis of (17S,20S)-3β-(t-butyldimethylsilyloxy)-17,20-methan-5-pregnene-21-carbaldehyde (5)*

To a solution of compound 4 (0.85 g, 1.91 mmol) in dry DCM (100 mL) was added at 0 °C Dess-Martin periodinane (1.62 g, 3.82 mmol) and the reaction mixture was stirred at room temperature for 1.5 hours. After completion of the reaction (checked by TLC), a mixture of saturated aqueous $NaHCO_3$ and 10% aq. $Na_2S_2O_3 \cdot 5H_2O$ (1:1) was added and the reaction mixture was stirred for 30 min. The reaction mixture was extracted with $Et_2O$ (30 mL x 3 times) and the combined organic layers washed with saturated aqueous $NaHCO_3$ and brine, dried over $Na_2SO_4$ and the solvent was removed *in vacuo*. The residue was purified by FCC (petroleum ether 40-60 °C/EtOAc 98:2→96:4) to afford compound 5 (0.72 g, yield 85%) as a white crystalline solid. Mp: 114–116 °C; $[\alpha]_D^{25}$= −21° (c= 0.0056 g/mL, $CHCl_3$); $R_f$: 0.78 (petroleum ether 40–60 °C/EtOAc 80:20); $^1H$ NMR (600 MHz, $CDCl_3$): $\delta$ 0.05 (s, 6H, $Si(CH_3)_2$), 0.80 (s, 3H), 0.88 (s, 9H, $C(CH_3)_3$), 0.92–0.99 (m, 1H), 1.00 (s, 3H), 1.04–2.30 (m, 21H), 3.42–3.52 (m, 1H, 3*α*-H), 5.33 (d, *J*= 5.1 Hz, 1H, 6-H), 9.08 (d, *J* = 6.2 Hz, 1H, CHO); $^{13}C$ NMR (75 MHz, $CDCl_3$): $\delta$ -4.4, 17.1, 18.4, 19.6, 20.5, 21.1, 25.3, 26.0, 29.6, 31.0, 32.0, 32.1, 32.6, 33.1, 36.8, 37.5, 42.3, 42.9, 43.8, 50.4, 53.9, 72.6, 120.9, 141.7, 202.0; APCI-HRMS: m/z calculated for $C_{28}H_{47}O_2Si$ $[M+H]^+$ 443.3340, found 443.3339.

*Synthesis of 3-((2'R,3S,17S)-3-((t-butyldimethylsilyl) oxy)-5-androstene-17,1'-cyclopropan]-2'-yl) acrylonitrile (6)*

To a suspension of NaH 60% in mineral oil (30 mg, 0.70 mmol) in dry THF was added at 0 °C diethyl(cyanomethyl)phosphonate (0.1 mL, 0.64 mmol) and the reaction mixture was stirred at room temperature for 30 minutes. Subsequently, a solution of compound **5** (70 mg, 0.16 mmol) in dry THF (0.1 M) was added at 0°C and the reaction mixture was stirred at the same temperature for 30 minutes. After completion of the reaction, the reaction mixture was quenched with saturated aqueous $NH_4Cl$ and extracted with EtOAc. The organic layer was washed with brine, dried over $Na_2SO_4$ and the solvent was removed *in vacuo*. The residue was purified by FCC (petroleum ether 40-60 °C/EtOAc 99:1→98:2) to afford compound **6-*Z*** (23 mg) and **6-*E***, (36 mg) in 84% overall yield as white crystalline solids.

*(Z)-3-((2'R,3S,17S)-3-((t-butyldimethylsilyl)oxy)-5-androstene-17,1'-cyclopropan]-2'-yl) acrylonitrile (6-Z)*

Mp: 130-132 °C; $[\alpha]_D^{25}$= -117.6° (c= 0.0034 g/mL, $CHCl_3$); $R_f$: 0.67, petroleum ether 40-60 °C/EtOAc 90:10; $^1H$ NMR (600 MHz, $CDCl_3$): $\delta$ 0.06 (s, 6H, $Si(CH_3)_2$), 0.54 (t, *J* = 5.0 Hz, 1H), 0.81 (s, 3H), 0.89 (m, 9H), 1.01 (s, 3H), 1.12-2.31(m, 20H), 3.49-3.53 (m, 1H, 3*α*-H) 5.32 (d, *J* = 4.9 Hz, 1H, 6-H), 5.19 (d, *J* = 10.8 Hz, 1H), 6.03 (t, *J* = 10.8 Hz, 1H); $^{13}C$ NMR (150 MHz, $CDCl_3$): $\delta$ -4.4, 16.7, 18.4, 19.6, 20.5, 22.2, 23.3, 25.2, 26.1, 29.9, 32.1, 32.2, 32.6, 33.7, 36.9, 37.6, 42.1, 42.3, 42.9, 50.5, 54.5, 72.7, 95.9, 117.2, 120.9, 141.8, 158.0; APCI-HRMS: m/z calculated for $C_{30}H_{48}NOSi$ $[M+H]^+$ 466.3500, found 466.3496.

*(E)-3-((2'R,3S,17S)-3-((t-butyldimethylsilyl)oxy)-5-androstene-17,1'-cyclopropan]-2'-yl) acrylonitrile (6-E)*

Mp: 193-196 °C; $[\alpha]_D^{25}$= + 40° (c= 0.0015 g/mL, $CHCl_3$); $R_f$: 0.64, petroleum ether 40-60 °C/EtOAc 90:10; $^1H$ NMR (600 MHz, $CDCl_3$): $\delta$ 0.06 (s, 6H, $Si(CH_3)_2$), 0.53 (t, *J* = 4.9 Hz, 1H), 0.75 (s, 3H), 0.89 (m, 9H), 1.00 (s, 3H), 1.12-2.28 (m, 20H), 3.49-3.53 (m, 1H, 3*α*-H) 5.30 ( *J* = 3.0 Hz, 1H, 6-H), 5.31 (d, *J* = 16.0 Hz, 1H), 6.30 (dd, *J* = 16.0, 10.2 Hz); $^{13}C$ NMR (150 MHz, $CDCl_3$): $\delta$ -4.4, 16.7, 18.4, 19.6, 20.5, 22.9, 23.1, 25.2, 26.1, 29.9, 32, 32.2, 32.6, 33.3, 36.8, 37.5, 42.2, 42.3, 42.9, 50.4, 54.6, 72.6, 96.2, 118.3, 121, 141.7, 159; APCI-HRMS: m/z calculated for $C_{30}H_{48}NOSi$ $[M+H]^+$ 466.3500, found 466.3497.

*Synthesis of (E)-3-((2'R,3S,17S)-3-(hydroxy)-5-androstene-17,1'-cyclopropan]-2'-yl) acrylonitrile (ENT-A011)*

To a solution of compound **6-*E*** (34 mg, 0.07 mmol) in anhydrous DCM (2.3 mL) HF·pyridine (0.1 mL) was added at 0 °C and the reaction mixture was stirred at room temperature for 2 hours. The reaction was quenched with water at 0 °C, and the resulting mixture was extracted with DCM. The organic layer was washed with brine, dried over $Na_2SO_4$ and the solvent was removed *in vacuo*. The residue was purified by FCC (Hexane/EtOAc 90:10→80:20) to afford compound **ENT-A011** (25 mg, yield quantitative) as a white crystalline solid. Mp: 180-183 °C; $[\alpha]_D^{25}$= +28° (c= 0.0029 g/mL, $CHCl_3$); $R_f$: 0.17, Hexane/EtOAc 80:20; $^1H$ NMR (600 MHz, $CDCl_3$): $\delta$ 0.53 (t, $J$ = 4.9 Hz, 1H), 0.76 (s, 3H), 0.83-0.98 (m, 2H), 1.01 (s, 3H), 1.12-2.28 (m, 19H), 3.49-3.53 (m, 1H, 3$\alpha$-H), 5.31 (d, $J$ = 16.0 Hz, 1H), 5.36 (m, 1H, 6-H), 6.30 (dd, $J$ = 16.0, 10.2 Hz, 1H); $^{13}C$ NMR (75 MHz, $CDCl_3$): $\delta$ 16.7, 19.5, 20.6, 22.9, 23.1, 25.1, 29.8, 31.7, 32.0, 32.5, 33.3, 36.7, 37.4, 42.1, 42.2, 42.3, 50.3, 54.5, 71.8, 96.2, 118.3, 121.6, 140.9, 159.1; APCI-HRMS: m/z calculated for $C_{24}H_{34}NO$ $[M+H]^+$ 352.2635, found 352.2628. HPLC: $t_R$ = 6.32 min, λ = 237 nm, purity = 99.09%.

*Synthesis of (Z)-3-((2'R,3S,17S)-3-(hydroxy)-5-androstene-17,1'-cyclopropan]-2'-yl)acrylonitrile (ENT-A012)*

To a solution of compound **6-*Z*** (22 mg, 0.04 mmol) in anhydrous DCM (1.5 mL) HF·pyridine (70 µL) was added at 0 °C and the reaction mixture was stirred at 25 °C for 2 hours. The reaction was quenched with water at 0 °C and the resulting mixture was extracted with DCM. The organic layer was washed with brine, dried over $Na_2SO_4$ and the solvent was removed in vacuo. The residue was purified by FCC (Hexane/EtOAc 90:10→80:20) to afford compound **ENT-A012** (13 mg, yield quantitative) as a white crystalline solid. Mp: 197-203 °C; $[\alpha]_D^{25}$= -157° (c= 0.0049 g/mL, $CHCl_3$); $R_f$: 0.05, Hexane/EtOAc 90:10; $^1H$ NMR (600 MHz, $CDCl_3$): 0.54 (t, $J$ = 5.0 Hz, 1H), 0.82 (s, 3H), 0.85-0.98 (m, 2H), 1.02 (s, 3H), 1.12-2.28 (m, 19H), 3.49-3.53 (m, 1H, 3α-H) 5.36 (d, $J$ = 5.1 Hz, 1H, 6-H), 5.19 (d, $J$ = 10.8 Hz, 1H), 6.03 (t, $J$ = 10.8 Hz, 1H); $^{13}C$ NMR (75 MHz, $CDCl_3$): δ 16.8, 19.6, 20.5, 22.3, 23.3, 25.2, 29.8, 31.7, 32.0, 32.6, 33.2, 36.7, 37.4, 42.1 42.2, 42.3, 50.4, 54.4, 71.9, 95.9, 117.3, 121.5, 141.0, 158.1; APCI-HRMS: m/z calculated for $C_{24}H_{34}NO$ $[M+H]^+$ 352.2635 found 352.2632. HPLC: $t_R$ = 6.32 min, λ = 237 nm, purity = 98.47%.

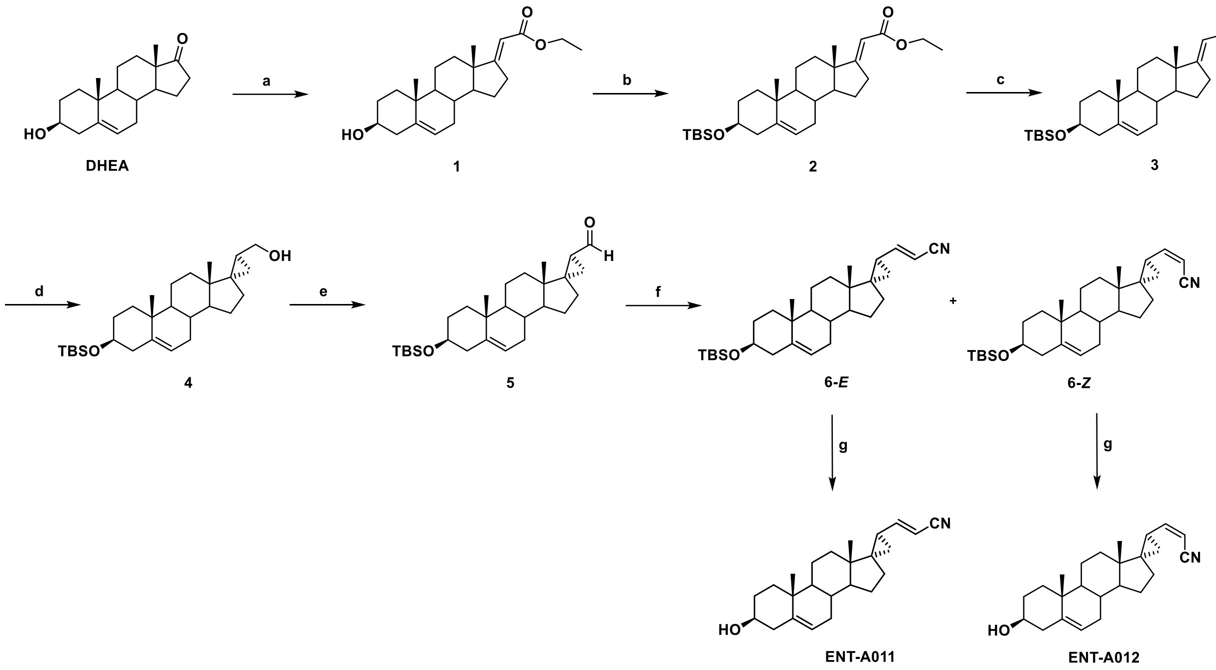


**Scheme 1. Synthesis of ENT-A011 and ENT-A012.** Reagents and conditions: (a) $(CH_3CH_2O)_2P(O)CH_2C(O)OCH_2CH_3$, EtONa, THF/EtOH 1:1, reflux, overnight; (b) TBSCl, Imidazole, $I_2$, THF, 0 °C to 25 °C, overnight; (c) DIBAL-H, $CH_2Cl_2$, −78 °C, 2.5 h; (d) $CH_2I_2$, $Et_2Zn$, −78 °C to 25 °C, 1 h; (e) DMP,$CH_2Cl_2$, 0 °C to 25 °C, 1.5 h; (f) $(CH_3CH_2O)_2P(O)CH_2CN$, NaH, THF, 0 °C to 25 °C, 0.5 h; (g) HF·Pyridine, $CH_2Cl_2$, 0 °C to 25 °C, 1 h.

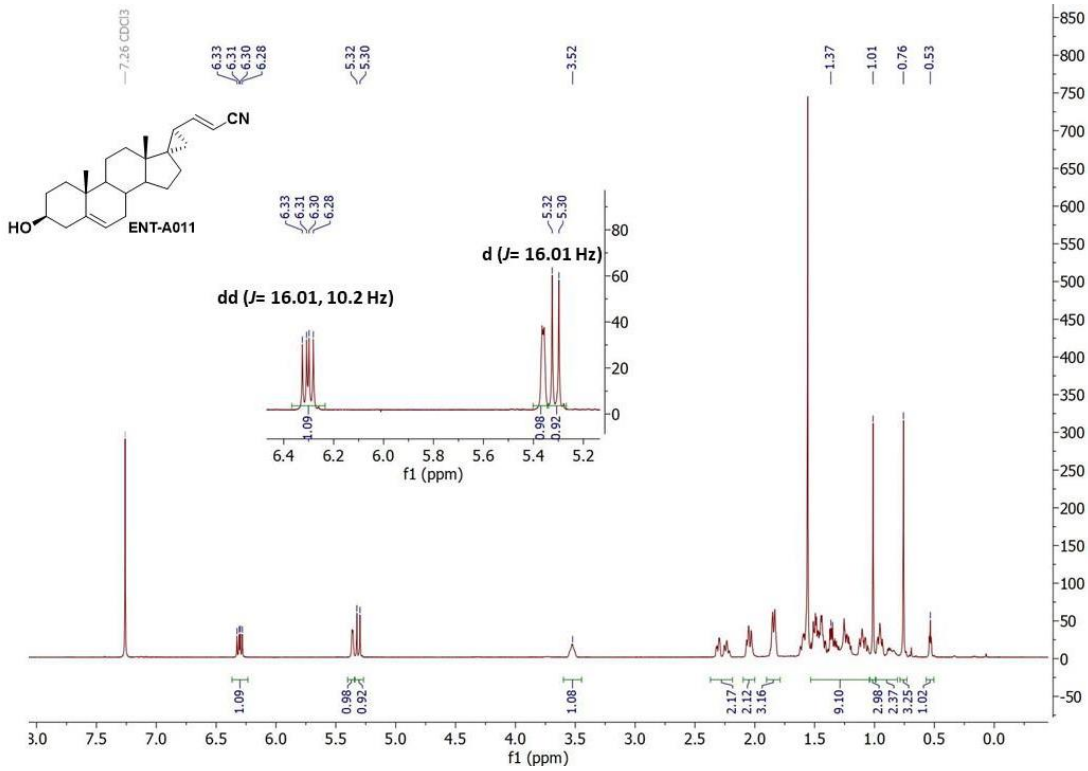


Figure S1. $^{1}$H-NMR spectrum of ENT-A011 in $CDCl_3$.

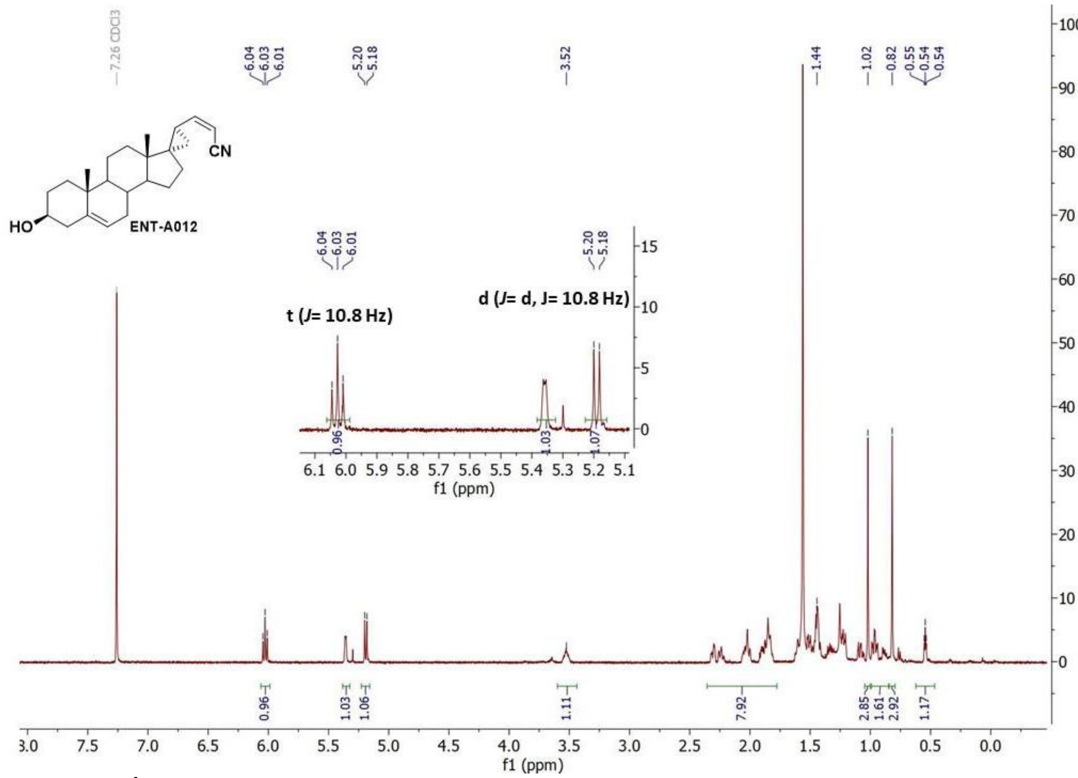


Figure S2. $^{1}$H-NMR spectrum of ENT-A012 in $CDCl_3$.

**Area % Report**

Data File:
C:\ChromQuest\Enterprise\Projects\Default\Method\TC\Alessia\ENT-A011_MeOH-H2O-01%HCOOH_25min-237nm.dat
Method:
C:\ChromQuest\Enterprise\Projects\Default\Method\TC\Alessia\Gradient_MeOH_H2O_0.1%HCOOH_25min_237nm_95-5to100_2.met
Acquired: 11/1/2021 2:33:08 PM
Printed: 11/1/2021 3:01:00 PM

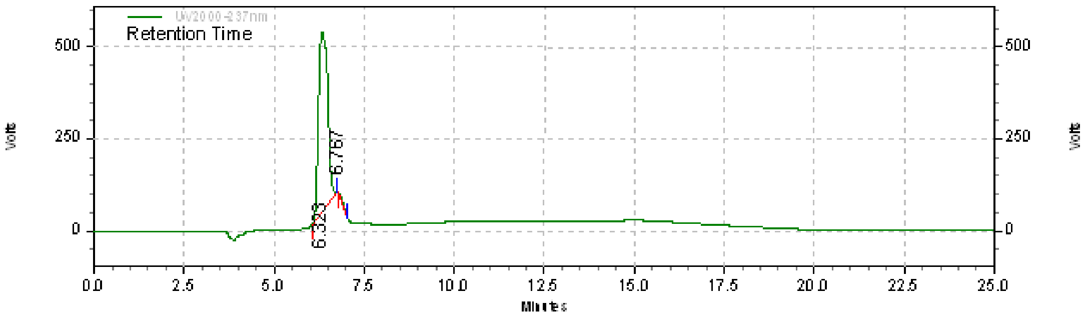


**UV2000-237nm Results (System (11/1/2021 3:00:57 PM) (Reprocessed))**

| Retention Time | Area | Area % | Height | Height % |
|---|---|---|---|---|
| 6.323 | 8678720 | 99.09 | 489972 | 100.00 |
| 6.767 | 79379 | 0.91 | 0 | 0.00 |
| Totals | 8758099 | 100.00 | 489972 | 100.00 |

**Figure S3. HPLC chromatogram of compound ENT-A011.** The purity of ENT-A011 was determined by high-performance liquid chromatography (HPLC) using Nucelosil 100-5 C18 HD column, 5µm (4.6 x 250 mm), flow rate 1 mL/min, eluting with $H_2O$, 0.1% HCOOH – MeOH, 0.1% HCOOH gradient employing UV detection at 237 nm. $t_R$ = 6.32 min, purity = 99.09%.

**Area % Report**

Data File:
C:\ChromQuest\Enterprise\Projects\Default\Method\TC\Alessia\ENT-A012_MeOH-H2O-01%HCOOH_25min-237nm.dat
Method:
C:\ChromQuest\Enterprise\Projects\Default\Method\TC\Alessia\Gradient_MeOH_H2O_0.1%HCOOH_30min.met

Acquired: 11/1/2021 3:07:18 PM
Printed: 11/1/2021 3:42:50 PM

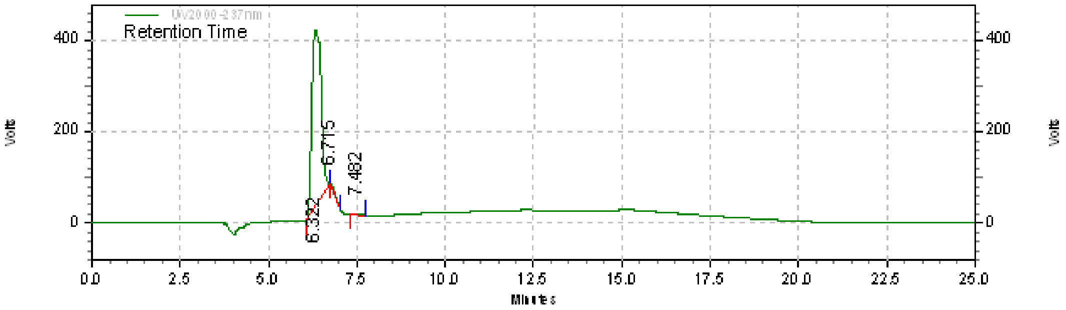


**UV2000-237nm Results (System (11/1/2021 3:42:43 PM) (Reprocessed))**

| Retention Time | Area | Area % | Height | Height % |
|---|---|---|---|---|
| 6.322 | 7009356 | 98.47 | 387982 | 99.62 |
| 6.715 | 84181 | 1.18 | 0 | 0.00 |
| 7.482 | 24607 | 0.35 | 1483 | 0.38 |
| Totals | 7118144 | 100.00 | 389465 | 100.00 |

**Figure S4**. **HPLC chromatogram of compound ENT-A012.** The purity of ENT-A012 was determined by high-performance liquid chromatography (HPLC) using Nucelosil 100-5 C18 HD column, 5µm (4.6 x 250 mm), flow rate 1 mL/min, eluting with $H_2O$, 0.1% HCOOH – MeOH, 0.1% HCOOH gradient employing UV detection at 237 nm. $t_R$ = 6.32 min, purity = 98.47%.

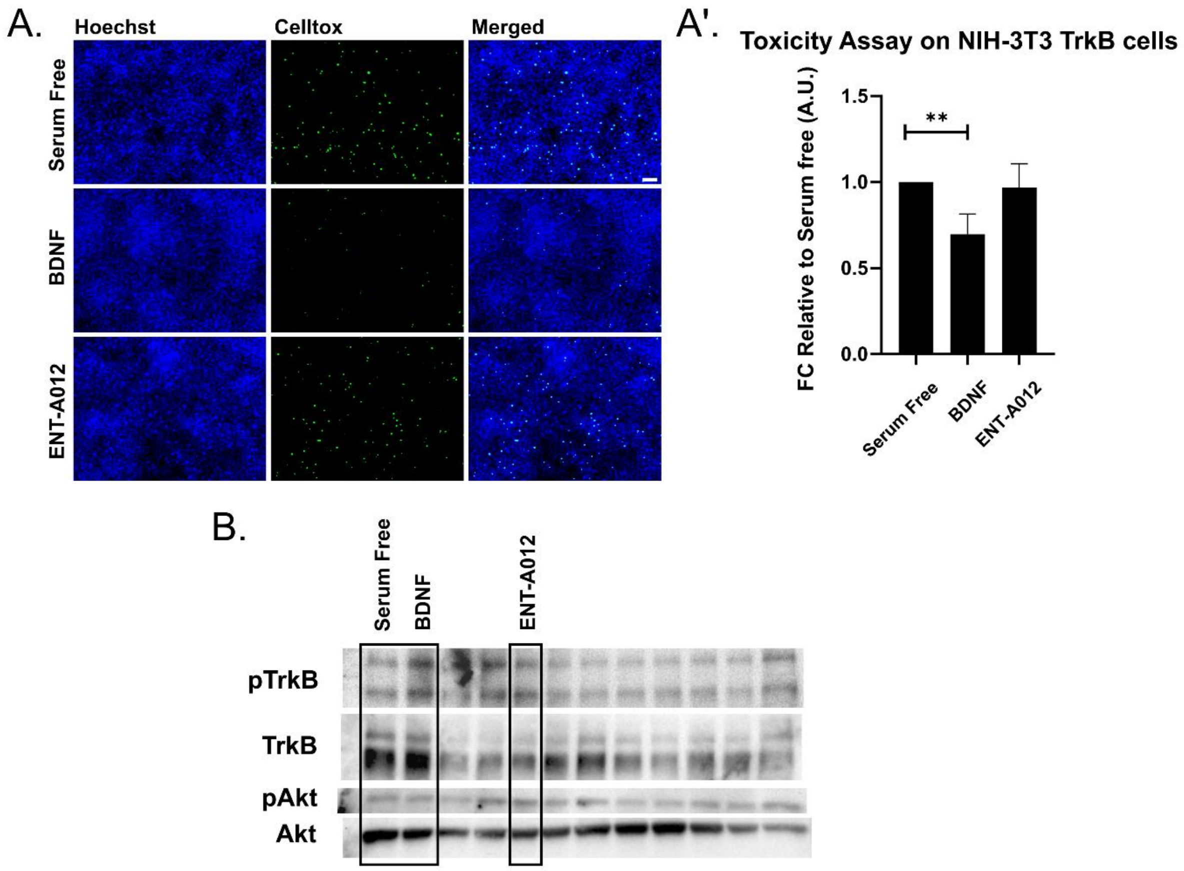


**Figure S5**. **ENT-A012 do not reduce cell death caused by serum deprivation in NIH-3T3 TrkB cells**. Representative images (A) and quantification of Toxicity Assay (A') in NIH-3T3 TrkB cells after treatment with BDNF or compound ENT-A012 for 24h in serum free conditions. N=13, error bars represent S.E.M., Student's t-test against Control; **<0.01. The compound ENT-A012 do not induce TrkB and its downstream target Akt phosphorylation after 20 minutes treatment in NIH-3T3 TrkB stable expressed cells, representative blot (B). Scalebar = 100µm

### Computational Study of ENT-A011 TrkB Docking

Computational docking studies were performed to investigate the mechanism of action of the compound ENT-A011. The extracellular domain of TrkA has been previously reported to be a drug target[3–5], while prior Saturation Transfer Difference Nuclear Magnetic Resonance (STD-NMR) experiments and molecular dynamics simulations have indicated that an analogue of ENT-A011, BNN27, binds at the interface of TrkA-D5 with NGF, thus bridging the heterodimer.[6] Specifically, two binding sites, site 1a and site 1b, found at the interfaces of the two proteins were proposed to be the most probable binding sites for BNN27.[6] Since there is high structural similarity between the TrkA and TrkB receptors, and ENT-A011 is an analogue of BNN27, the corresponding binding sites 1a and 1b of TrkB were identified in the present work and used for docking (Figure S6). The binding poses of the compound in the two sites show complementarity to the binding pockets. For site 1b, the docking pose has the 3β-hydroxyl group pointing towards the solvent region and the C17 steroid substituent facing towards the interface of TrkB-D5 and NT-4/5, thus providing a possible explanation for the selectivity of the substituent. The binding pose in site 1a is less solvent-exposed and preliminary molecular dynamics simulations showed that the compound bound in this pocket dissociates less readily than from site 1b (data not shown). Overall, the docking studies suggest two plausible interaction modes of ENT-A011 with TrkB that could lead to enhanced receptor activation.

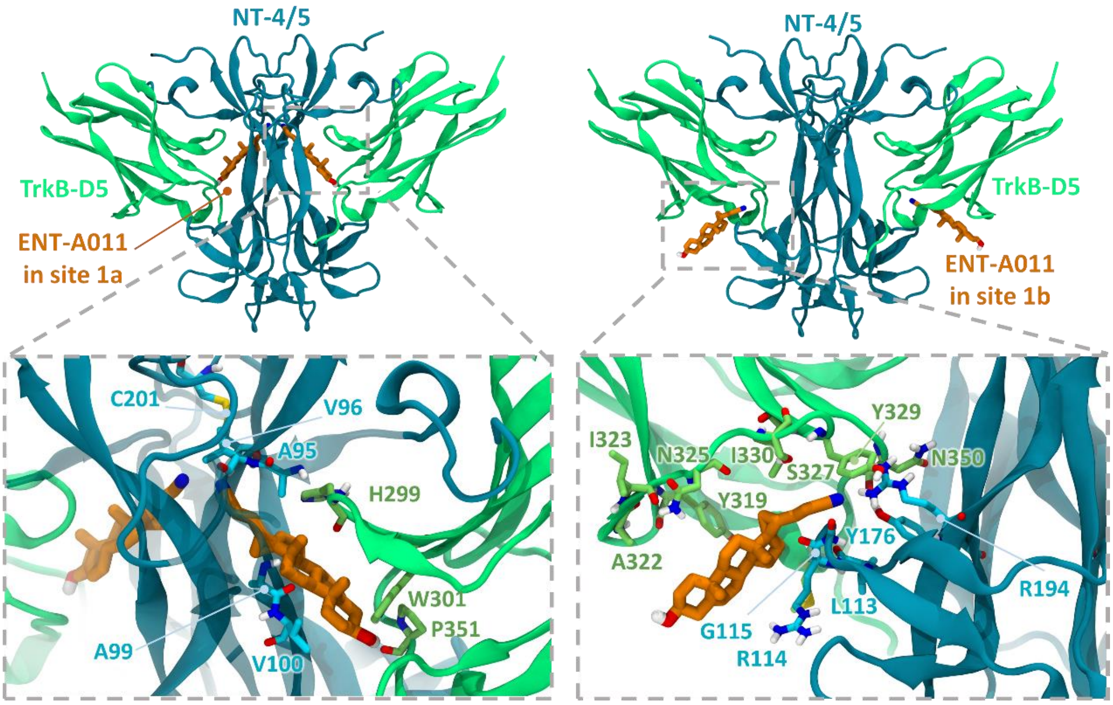


**Figure S6**: Docking poses of compound ENT-A011 in sites 1a and 1b at the two symmetry-related interfaces of the NT-4/5 -TrkB-D5 neurotrophin-receptor complex. The close-up views show residues lining the two sites. TrkB-D5 and NT-4/5 are shown, respectively, in green and blue cartoon representation with selected residues in stick representation colored by atom type. The ENT-A011 compound is shown in stick representation colored by atom type with orange carbons.

### ENT-A011 exhibits very slow depletion in human liver microsomes

The susceptibility of a test-compound to biotransformation is defined as metabolic stability. For the ranking of test compounds, several approaches can be followed including parent structure loss during metabolic reactions or their intrinsic clearance ($CL_{int}$) and *in vitro* half-life ($t_{1/2}$) values[1,2]. Herein, we chose the former.
Parent structure loss is classified as very slow (<5 %), slow (5-19 %), moderate (20-50 %), fast (50-80 %) or very fast (>80 %). Such categories have been defined according to set criteria, namely, high metabolism ($t_{1/2}$ value of <30 min), moderate metabolism (30 min < $t_{1/2}$ value of < 60 min) and low metabolism ($t_{1/2}$ value of >60min). ENT-A011 is very slowly depleted showing 93% residual of time zero at t=60 minutes. Thus, ENT-A011 may correspond to low or medium intrinsic clearance classification bands.
For humans, a low intrinsic clearance classification band is defined by an $CL_{int}$ <8.6 μL/min/mg protein, whereas a high intrinsic clearance classification band is defined by an $CL_{int}$ >47.0 μL/min/mg protein. Low clearance test-compounds are characterized by enhanced exposure, prolonged half-life and reduced doses, predicted as suitable for once-daily dosing.

**ENT-A011 shows weak-to-moderate CYP inhibition**

Human liver cytochrome P450 (CYP450) enzymes are crucial for xenobiotic biodegradation, metabolism, and toxicity as well as xenobiotic-host and/or xenobiotic-xenobiotic interactions. Upon linear velocity conditions *in vitro*, the depletion rate of ENT-A011 may be extrapolated to a. *in vivo* hepatic clearance, b. extraction ratio, and c. the effect of hepatic first-pass metabolism to total oral bioavailability. Biodegradation, metabolic, and toxicity liabilities can be identified early on and thus, inform structure-activity relationships (SAR).

For this, after the administration of ENT-A011 at 1 μM, the activity of CYP1A2, CYP2A6, CYP2B6, CYP2C9, CYP2C19, CYP2D6, and CYP3A4 (seven major human CYP450s) was assessed to determine a. the oxidative (CYP-mediated) metabolic stability profile in question and b. the enzyme metabolizing isoforms responsible (the test system consists of recombinant human CYP450 and CYP450 reductase; cytochrome $b_5$ may also be present).

Herein, no concentration-dependent effects are anticipated, as our model depicts direct interactions between ENT-A011 and CYP450 isoenzymes. CYP450 substrates are known to change enzyme conformation, disrupt enzyme structure and/or function and/or block the enzyme active site, altering enzyme[7,8]. First, the catalytic activity of CYP1A2, CYP2A6, CYP2B6, CYP2C9, CYP2C19, CYP2D6, and CYP3A4 was determined after the administration of ENT-A011 at 1 μM. Catalytic activity was assessed on the basis of the relative fluorescence of the enzymatic reaction product (Figure S6_A). ENT-A011 did not show strong effects on the enzyme (catalytic) activity of the seven major CYP450-isozymes tested herein.

Next, CYP450 (%) inhibition was determined. For the CYP450 isoenzymes tested, no product inhibition or mechanism-based inactivation was obtained. No solubility issues were reported for ENT-A011 (new chemical entities with poor solubility may result in artificially low CYP450 inhibition and thus, potential drug-drug interaction toxicities may escape our attention). CYP450 enzyme inhibition may result in unexpectedly high exposure of co-administered xenobiotics and hence, increase the risk for adverse effects. As depicted in Figure S6_B, ENT-A011 exhibits weak inhibition for CYP2D6, while it is a moderate inhibitor for CYP2B6, CYP2C9, and CYP2C19. For all CYP450 isoforms tested, we also assessed CYP450 (%) metabolic activity, expressed as residual % of time zero. Our findings are depicted in Figure S6_C, according to which ENT-A011 was overall metabolically stable (from t=0 to t=60 minutes).

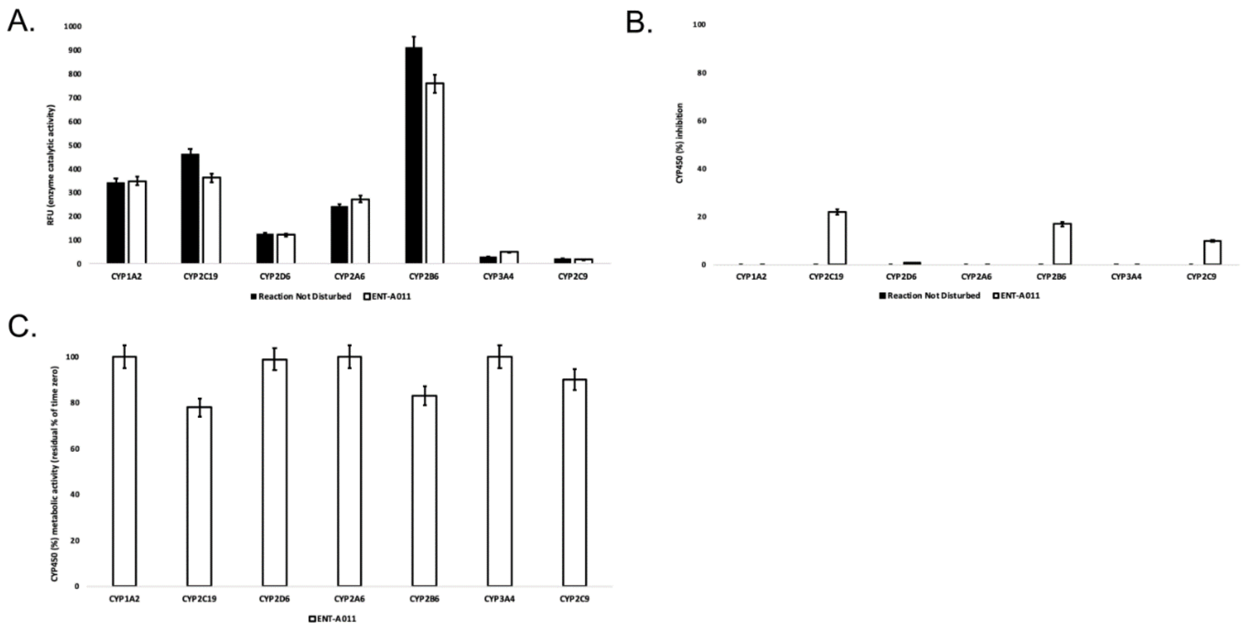


**Figure S7**. A. Enzyme (catalytic) activity of the CYP1A2, CYP2A6, CYP2B6, CYP2C9, CYP2C19, CYP2D6, and CYP33A4 isoenzymes, following ENT-A011 administration at 1 μM (t=60 minutes). RFU: relative fluorescence units. Reaction Not Disturbed: reaction without ENT-A011. B. CYP450 (%) inhibition per isozyme tested after the administration of ENT-A011 at 1 μM (t=60 minutes). C. CYP450 (%) metabolic activity of CYP1A2, CYP2A6, CYP2B6, CYP2C9, CYP2C19, CYP2D6, and CYP33A4 isozymes, following ENT-A011 administration at 1 μM (t=60 minutes).

Herein, we performed an isozyme-specific CYP450-study to delineate ENT-A011 interactions with each of the seven major CYP450s that account for xenobiotic biodegradation, metabolism, and toxicity as well as xenobiotic-xenobiotic and/or xenobiotic-host interactions.

ENT-A011 acts as a weak-to-moderate CYP450 enzyme inhibitor, yet showing no biodegradation, liver metabolism or safety issues.

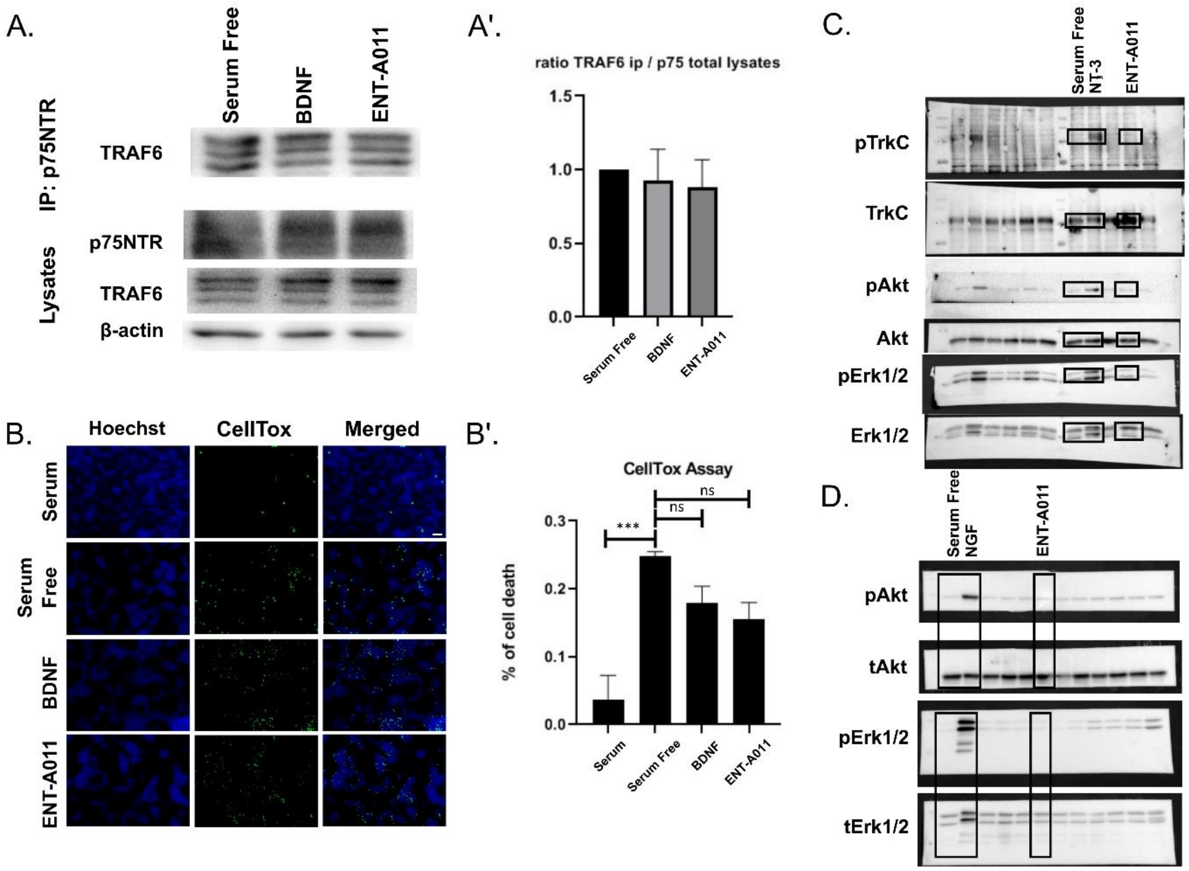


**Figure S8**. **ENT-A011 does not activate the p75 pathway.** A. HEK293T cells were co-transfected with the plasmid cDNAs of p75NTR and TRAF6. Transfectants were exposed for 20 min to BDNF (500ng/ml) & the tested compound ENT-A011 (1μM) and lysates were immunoprecipitated with p75NTR-specific antibodies and then immunoblotted with antibodies against TRAF6. Total lysates were analysed for p75NTR, TRAF6 and actin expression by immunobloting. A'. Quantification shows that ENT-A011 does not activate p75NTR *(one way ANOVA, no significance, Mean±SEM of triplicate measurements)*. B. HEK cells transfected with p75NTR were starved from serum and treated with ENT-A011 (1μM) or BDNF (500ng/ml) for 24hrs and subsequently subjected to CellTox assay. B'. Quantification shows that there is no significant difference between ENT-A011 treated group and negative control, Serum free. C. NIH-3T3 TrkC stable transfected cells were treated with Neurotrophin-3 (NT-3) or ENT-A011 for 20min and the lysates were immunoblotted with antibodies against pTrkC, TrkC, pAkt, Akt, pErk1/2, Erk1/2. The compound ENT-A011 did not phosphorylate TrkC neither activate the downstream pathway D. PC12 cells were treated with NGF or ENT-A011 for 20 min, lysates were immunoblotted with pAkt, Akt, pErk1/2, Erk1/2. The TrkA signaling was not activated by the compound ENT-A011 in contrast to NGF. Scalebar = 100μm
